# Pseudo-mutant p53 as a targetable phenotype of *DNMT3A*-mutated pre-leukemia

**DOI:** 10.1101/2021.05.30.446347

**Authors:** Amos Tuval, Yardena Brilon, Hadas Azogy, Yoni Moshkovitz, Tamir Biezuner, Dena Leshkowitz, Tomer M Salame, Mark D Minden, Perry Tal, Varda Rotter, Moshe Oren, Nathali Kaushansky, Liran I Shlush

## Abstract

Pre-leukemic clones carrying *DNMT3A* mutations have a selective advantage and an inherent chemo-resistance, however the basis for this phenotype has not been fully elucidated.

Mutations affecting the gene *TP53* occur in pre-leukemic hematopoietic stem/progenitor cells (preL-HSPCs) and lead to chemo-resistance. Many of these mutations cause a conformational change and some of them were shown to enhance self-renewal capacity of preL-HSPCs.

Intriguingly, a misfolded p53 was described in AML blasts that do not harbor mutations in *TP53*, emphasizing the dynamic equilibrium between a wild-type (WT) and a “pseudomutant” conformations of p53.

By combining single cell analyses and p53 conformation-specific monoclonal antibodies we studied preL-HSPCs from primary human *DNMT3A* AML samples. We found that while leukemic blasts express mainly the WT conformation, in preL-HSPCs the pseudomutant conformation is the dominant. HSPCs from non-leukemic samples expressed both conformations to a similar extent.

Treatment with a short peptide that can shift the dynamic equilibrium favoring the WT conformation of p53, specifically eliminated preL-HSPCs that had dysfunctional canonical p53 pathway activity as reflected by single cell RNA sequencing.

Our observations shed light upon a possible targetable p53 dysfunction in human preL-HSPCs carrying *DNMT3A* mutations. This opens new avenues for leukemia prevention.

## Introduction

Although Acute Myeloid Leukemia (AML) is preceded by clonal hematopoiesis (CH), most CH clones are not “pre-leukemic” and do not transform to AML^1^. When referring to the most common mutated gene in CH, *DNMT3A*, the main discriminative characteristics between pre-AML and CH are larger clones and more accompanying mutations, reflecting the selective advantage of these clones^2^. Experimental and clinical studies have demonstrated that *DNMT3A* mutated pre-leukemic clones have also an inherent chemo-resistance and the ability to reconstitute the bone marrow following AML chemotherapeutic treatments^3–5^. The mechanisms behind this pre-AML phenotype remain unclear.

Mutations affecting *TP53* occur during the pre-leukemic stage of AML^6^. Some of these mutations cause a conformational change^7^ that can be detected using conformation-specific monoclonal antibodies^8^. In a heterozygous state, the mutant protein can have a dominant negative effect over the wild type (WT) protein that leads to chemoresistance and p53 dysfunction as reflected by a reduced expression of downstream target genes of p53^9^. Moreover, *TP53* mutations can enhance the self-renewal capacity of murine HSPCs^10^.

Interestingly, a dynamic equilibrium was described between the WT and the mutant conformations of p53. The WT protein can acquire a “pseudo-mutant” conformation rendering it dysfunctional, with a reduced transcriptional activity^11^. The pseudo-mutant conformation was found in *TP53* WT AML blasts^12^, and was correlated with growth factor stimulation^13^.

We hypothesized that in *DNMT3A*—mutated AML, p53 conformational changes occur during an early evolutionary stage, in preL-HSPCs. Furthermore, we decided to test the influence of a short peptide that stabilizes the WT conformation of p53 and restores its transcriptional activity, on the engraftment capacity of *DNMT3A* mutated preL-HSPCs in immuno-deficient mice.

## Methods

### Samples

Primary samples were received from Princess Margaret Cancer Centre, University Health Network (UHN), Canada (UHN IRB protocol 01-0573) and from Rambam Health Care Campus, Israel (IRB protocol #283-1). Clinical characteristics of each sample are presented in supplemental Table 1.

### Pseudo mutant p53 evaluation by mass cytometry

We used a panel of metal-conjugated monoclonal antibodies targeting surface markers (supplemental Table 2). The two conformations of p53 were identified using p53 conformation-specific monoclonal antibodies that were conjugated to heavy metals: pAb1620 (for the WT) and pAb240 (for the mutant) (see supplemental information).

Phenotypic preL-HSPCs were defined as cells that are negative for CD33, CD15, CD11b, CD19, CD79b, CD3, CD16 and CD45RA and are CD34 positive. When identified in primary AML samples, a portion of the cells with this immune-phenotype were shown to be pre-leukemic (harboring only *DNMT3A* mutation, without *NPM1* mutation)^3^. Leukemic blasts from the AML samples were gated according to the original flow cytometry immune-phenotype of each sample.

### Xenotransplantation assays and *in vivo* pharmacologic treatment

All experiments were performed in accordance with institutional guidelines approved by the Weizmann Institute of Science Animal Care Committee (11790319-2) and as described previously^3^. Primary CD3 depleted AML samples were injected to immune-deficient mice (see supplemental information). This mouse model exposes the rare preL-HSPCs that reside in the injected pool^3^. Five weeks after the injection of human cells, when engraftment was established, we treated some of the mice for two weeks with a short peptide. Other mice were treated with a scrambled, control peptide. Mice were sacrificed on day 56. We evaluated human engraftment by flow cytometry and sorted the main sub-populations for deep targeted DNA sequencing according to the mutations of the original injected pool, thus identifying their stem cell of origin (Figure 2A) (see supplemental information).

### Single cell RNA sequencing (scRNA-seq) of engrafting cells

Bone marrows of sacrificed mice were extracted and human cells were separated according to expression of human CD45 (EasySep^™^, StemCell Technologies, Vancouver, Canada). Libraries were prepared using 10X Genomics technology (see supplemental information).

### Statistical analyses

Comparisons between two groups, were performed using the two-tailed, non-paired, nonparametric Wilcoxon rank sum test with 95% confidence interval and continuity correction.

## Results and discussion

We analyzed p53 conformations by mass cytometry in nine primary *DNMT3A*-mutated, AML samples, a sample of healthy cord blood and two samples of mobilized PBMCs of *DNMT3A*^R882H^ CH (with a high variant allele frequency, VAF (supplemental Table 1)). All adult samples were deeply sequenced to verify that they do not carry *TP53* mutations even at low VAFs.

At single cell level, we calculated the ratio between the intensity of the pseudo-mutant conformation and that of the WT conformation of p53. We termed this ratio the “pseudo-mutant to WT conformation ratio”, PM/WT-CR. We found that, although there is wide variability with regard to the expression of the two conformations of p53, leukemic blasts express mainly the WT conformation (with a median PM/WT-CR of 0.53). Phenotypic preL-HSPCs were found to express mainly the pseudo-mutant conformation of p53 (a median PM/WT-CR of 3.06) (Figure 1A-B). HSPCs from cord blood and from *DNMT3A*^R882H^ CH (non-leukemic samples) were found to express both conformations to a similar extent (Figure 1C, supplemental Figure 1).

**Figure 1:**
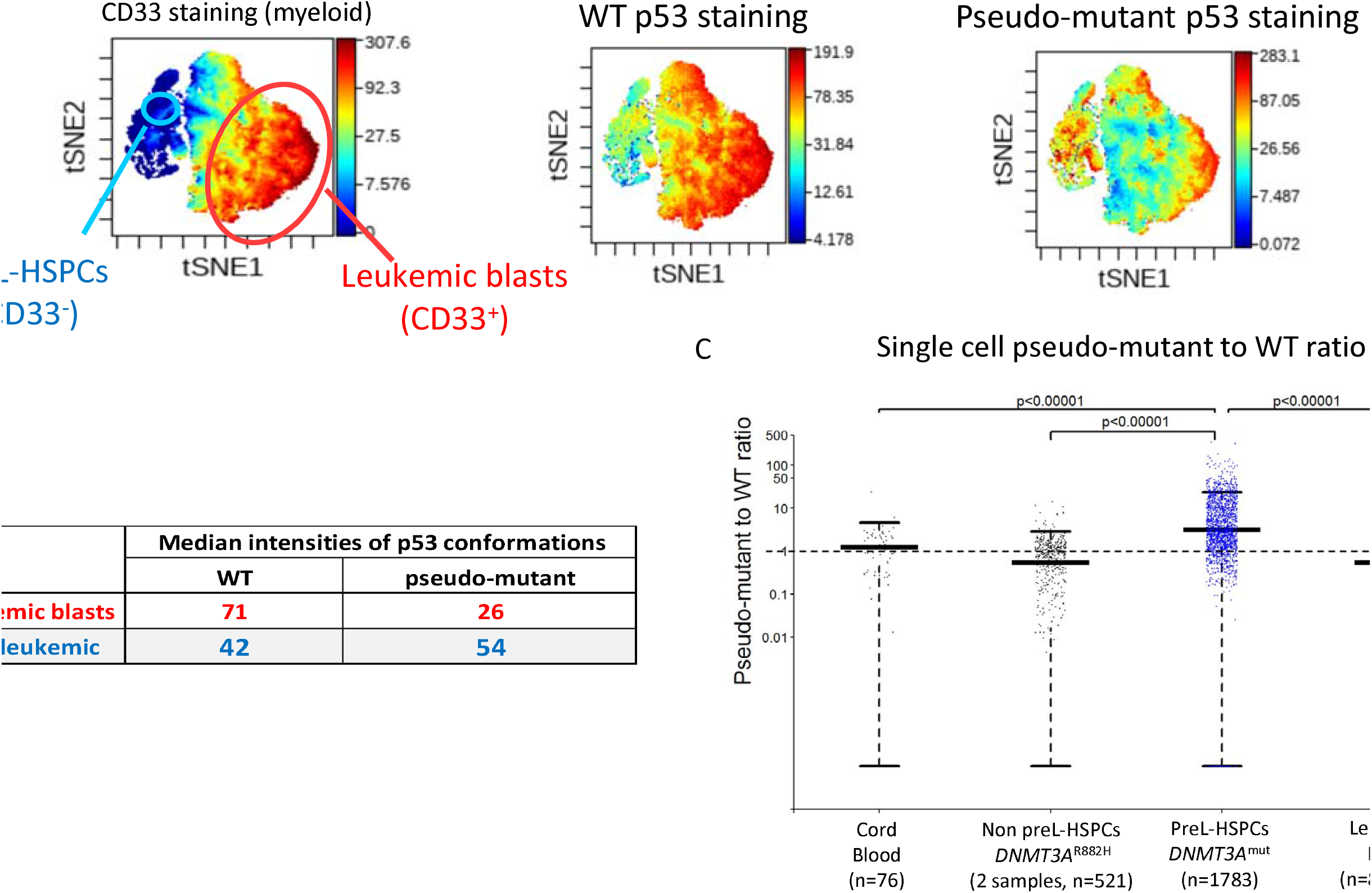
Mass cytometry of primary human samples. Mass cytometry (A) and median intensities of p53 conformations (B) in leukemic blasts (red) and in pre-leukemic HSPCs (blue) identified in the peripheral blood of a *DNMT3A*^R882H^, *NPM1c* AML patient. Using conformation-specific monoclonal antibodies we found that blasts express mainly the wild-type (WT) conformation of p53, while the dominant conformation in pre-leukemic cells is the pseudo-mutant conformation. (C) Ratio between the pseudo-mutant conformation and the WT conformation of p53 at the single cell level in different samples: HSPCs from a cord blood sample, HSPCs from 2 samples of *TP53* WT, *DNMT3A*^R882H^ CH, HSPCs from 9 *TP53* WT, *DNMT3A*-mutated, AML samples and blasts from the same AML samples. In preL-HSPCs (blue) the dominant conformation of p53 is the pseudo-mutant, with the highest ratio. This ratio was around 1 (dashed line) in HSPCs from non-leukemic samples (black). In the leukemic blasts (red), the dominant conformation is the WT with a median ratio of less than 1. n denotes the number of cells analyzed in each cohort. The median and 1.5 X interquartile range, are presented. The two-tailed, non-paired, nonparametric Wilcoxon rank sum test was used with 95% confidence interval and continuity correction.

**Figure 2:**
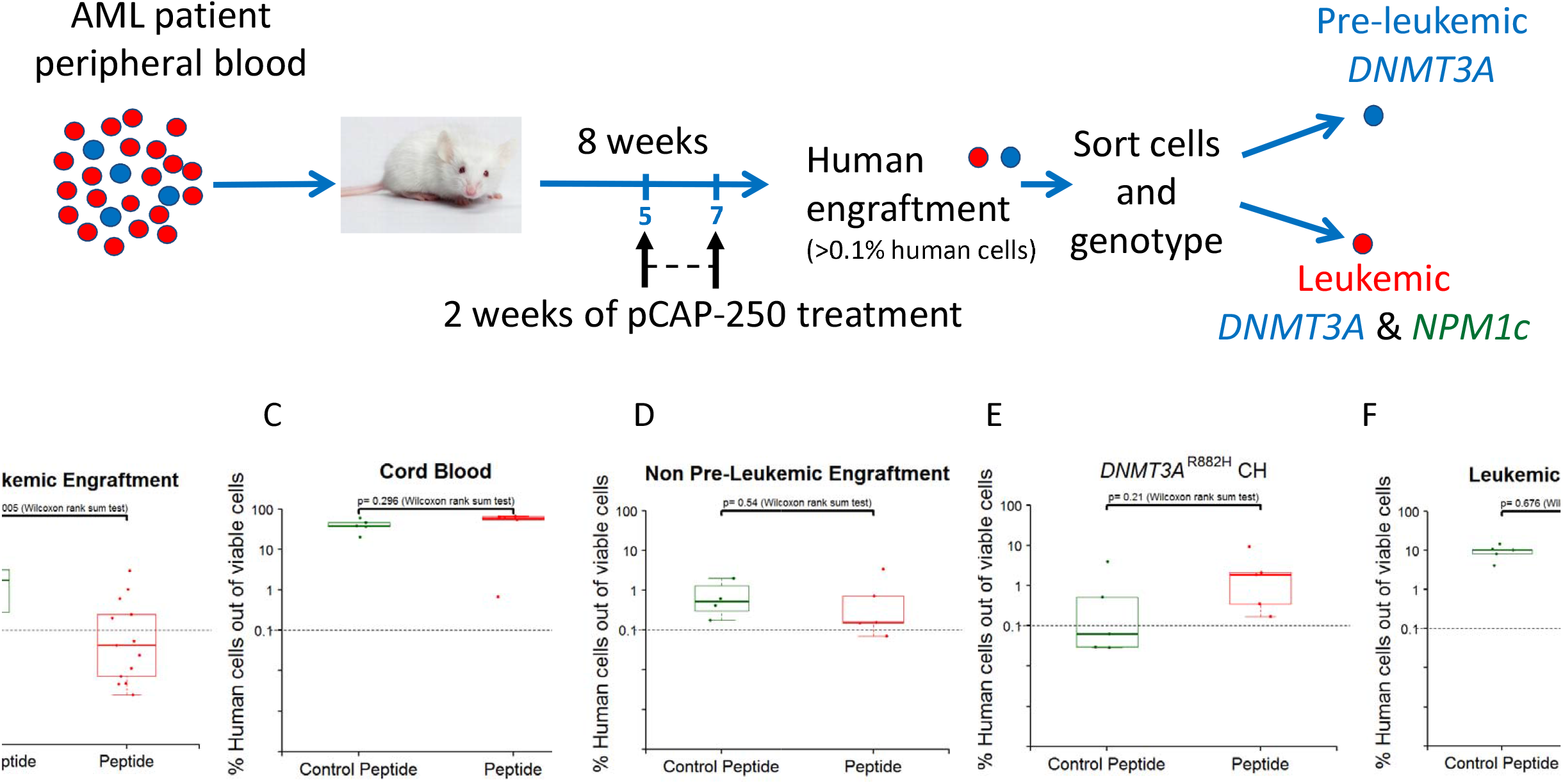
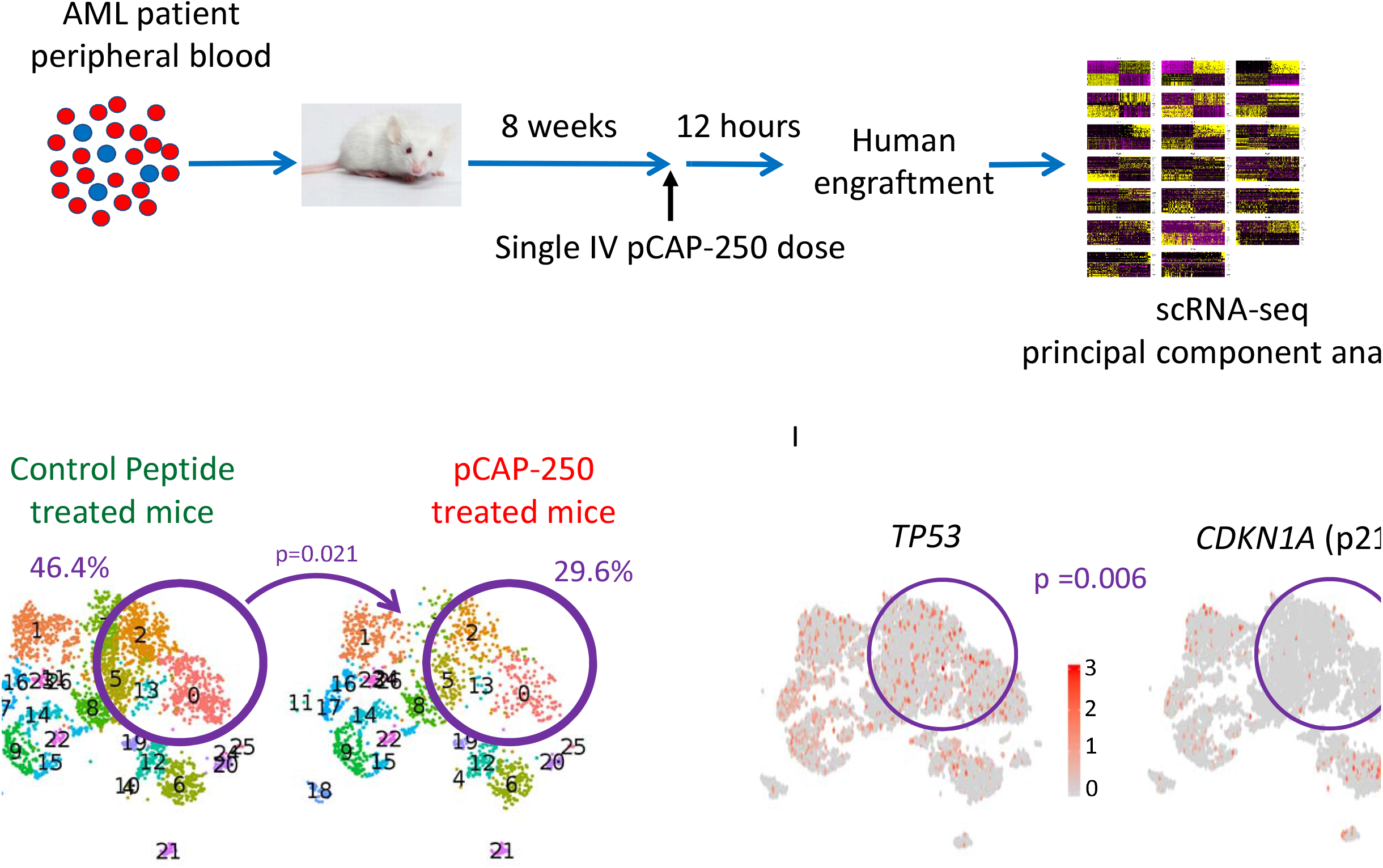
Xenotransplantation assays and *in vivo* pharmacologic treatment. (A) Schematic of the experimental setup. The pool of PBMCs of AML patients contains mainly blasts (red circles) but also rare preL-HSPCs (blue circles). Sorting and sequencing the engrafting cells allowed us to determine whether they originate from preL-HSPCs (having only a *DNMT3A* mutation) or from leukemic stem cells (having both *DNMT3A* and *NPM1* mutations). This experimental model enabled us to explore the vulnerability of the different cells to p53-directed treatment. (B) -(F) Engraftment of the different injected samples as assessed by flow cytometry according to treatment cohort: control peptide (green) and pCAP-250 treated (peptide, red). The origin of the engrafting cells was determined by sequencing (see supplemental Figure 2). Each dot represents a mouse. The dashed line represents the threshold for engraftment (presence of 0.1% of human cells out of all cells extracted from the bone marrow). Box plot centers, hinges and whiskers represent the median, first and third quartiles and 1.5 X interquartile range, respectively. Boxes are drawn with widths proportional to the square-roots of the number of observations in each cohort. All comparisons were performed using a two-tailed, non-paired, nonparametric Wilcoxon rank sum test with 95% confidence interval and continuity correction. (G) Schematic of the experimental setup for gene expression profiling of engrafting cells. (H) tSNE analyses of scRNA-seq data of engrafting cells. Each dot represents a cell. Both treatment cohorts contain the same sub-populations, however, some clusters (0,2,5,13 and 20 - circled in purple) decreased quantitatively following treatment with pCAP-250. This decrease from 46.4% to 29.6%, was found to be significantly dependent on treatment cohort (p-value 0.021, chi square test of independence with Yates continuity correction). (I) tSNE analyses of scRNA-seq data of engrafting cells with the expression of *TP53* (left) and of *CDKN1A* (p21, right). We calculated the difference between the logarithmic fold change of the expression of *CDKN1A* and that of *TP53* in pCAP-250 affected cells (clusters 0,2,5,13) and compared it to the same difference that was calculated for less affected cells (clusters 1,6-9,12,14-19,21-23). We found that affected cells had a significantly reduced expression of *CDKN1A* relative to their *TP53* expression, when compared to other clusters of cells (p = 0.006, Wilcoxon rank sum test).

We speculated that the high PM/WT-CR can contribute to the enhanced selective advantage of preL-HSPCs^10,14^ and that reverting this ratio might reduce their fitness. To achieve this, we used a short peptide (myr-RRHSTPHPD, named: pCAP-250) that can shift the balance between the WT and the mutant conformations of p53 by stabilizing the wild-type conformation. This peptide was shown to restore the downstream transcriptional activity of p53^15^. We tested the influence of pCAP-250 on the engraftment capacity of *DNMT3A, NPM1* mutated AML samples. One of the samples (sample #160005) gave rise to a multi-lineage graft that was composed of pre-leukemic cells harboring only the pre-leukemic *DNMT3A* mutation (supplemental Figure 2A-B). The engraftment capacity of these cells decreased significantly following treatment with pCAP-250 in three independent experiments (Figure 2B). pCAP-250 treatment did not affect the engraftment capacity of cord blood, non-preL-HSPCs from an AML sample, *DNMT3A*^R882H^ CH or of leukemic stem cells (Figure 2C-F, supplemental Figure 2C-E).

To conclude this part, we provided evidence that phenotypic preL-HSPCs derived from *DNMT3A* mutated AML samples have high levels of pseudo mutant p53 and are sensitive to p53-directed pharmacologic intervention. These data support the hypothesis that a high PM/WT-CR is associated with the increased fitness of preL-HSPCs.

Next, we explored whether these vulnerable pre-leukemic cells have p53 dysfunction by studying their transcriptional signature. To this end, we treated mice with a single dose of pCAP-250 eight weeks following injection of the same sample (#160005). We sacrificed the mice 12 hours later and performed scRNA-seq of the engrafting pre-leukemic cells (Figure 2G). Unbiased clustering of the engrafting cells revealed that while both treatment cohorts contained the same sub-populations, some clusters significantly decreased quantitatively following treatment with pCAP-250 (Figure 2H). The affected cells had a significantly reduced expression of p21, relative to their *TP53* expression level. Since p21 is the major canonical *TP53* regulated gene, this suggests that cells that were affected the most by the short peptide had a dysfunctional activity of the canonical pathway of p53 (Figure 2I).

In summary, our data demonstrate that the pseudo-mutant conformation switch of p53 occurs in the early stages of leukemia evolution and that it is partially lost following transformation to AML. While *DNMT3A* mutations might predispose for acquiring the pseudo-mutant phenotype, other factors may also influence the dynamic equilibrium between the two conformations. Previous studies have demonstrated that cytokines^13^, and inflammation^16^ can modulate this equilibrium. This might explain the heterogeneity among cells that belong to similar leukemia evolutionary stages.

Our results highlight that the pseudo-mutant phenotype is not merely a marker of preleukemia, rather it has functional consequences^17^. These cells probably have p53 dysfunction which can be targeted therapeutically. Of note, *DNMT3A* mutated primary AML do not present with a classical TP53-mutated phenotype (namely, a complex karyotype). This suggests that the pseudo-mutant p53 is not equivalent to a mutated p53. Future studies should explore the mechanisms that underlie the predisposition for this conformational switch in *DNMT3A*-mutated preL-HSPCs and its correlation with their increased fitness. It remains unclear whether this phenomenon can be generalized to other pre-leukemic mutations, however this offers new opportunities for AML prevention.

## Data Availability

The dataset generated and analyzed during the current study are available in the NCBI Sequence Read Archive (SRA; https://www.ncbi.nlm.nih.gov/sra/) under access numbers. Code is available on GitHub under https://github.com/ShlushLab

## Acknowledgments

The authors would like to thank Andrea Arruda for providing the clinical information of the primary samples and to Merav Kedmi from the department of life sciences core facilities at the Weizmann Institute of Science for technical guidance.

## Authorship Contributions

V.R., M.O. and L.S. initiated the project; A.T., N.K. and L.S. designed the research, and wrote the paper with input from other authors; A.T., N.K., H.A., Y.M., T.B. performed the research; A.T., Y.B., D.L. and T.M.S. analyzed the data; M.D.M. contributed clinical samples; P.T. contributed reagents and cell lines.

