## Supplemental information for "Pseudo-mutant p53 as a targetable phenotype of *DNMT3A*-mutated pre-leukemia"

**Supplemental information for Tuval A et al, Pseudo-mutant p53 as a targetable phenotype of *DNMT3A*-mutated pre-leukemia**

Samples

All samples were collected, Ficoll separated and viably frozen. CD3 cells were separated (EasySep™, StemCell Technologies, Vancouver, Canada) and expanded *in vivo* as previously described^1^. Mobilized peripheral blood mononuclear cells (PBMCs) and cord blood samples were also enriched for CD34^+^ cells (CD34 MicroBead Kit, Miltenyi Biotec, Bergisch Gladbach, Germany).

Mass cytometry

Maxpar® X8 Antibody Labeling Kit (Fluidigm, San Francisco, CA, USA) was used to conjugate antibodies to heavy metals.

Staining procedure was performed at room temperature (20°C). Briefly, 5x10^6^ Cells were incubated in Maxpar® PBS containing 1.25µM Cell-ID™ Cisplatin (Fluidigm, San Francisco, CA) at room temperature for 1 min, followed by incubation with antibodies directed at cell surface proteins (Supplemental table 2) for 30 min at room temperature in Maxpar® Cell Staining Buffer. Cells were washed fixed and permeabilized using the Maxpar® Nuclear Antigen Staining Buffer Set, and then incubated with antibodies directed at intra-nuclear proteins for 30 min at room temperature. Cells were washed and fixed in 4% Paraformaldehyde (Thermo Fisher Scientific, Waltham, MA) overnight at 4°C. Following this, cells were incubated in 125nM Cell-ID™ Intercalator-Ir solution (Fluidigm, San Francisco, CA), washed, resuspended in Maxpar® Water containing 1:10 EQ™ Four Element Calibration Beads and acquired via a CyTOF- Helios™ mass cytometer (Fluidigm, San Francisco, CA). Data were normalized and concatenated using CyTOF Software v.6.7.1014 and analyzed (i.e. gating and multidimensional analyses) using Cytobank (Cytobank Inc.;^2^). viSNE^3^ analyses were performed using all surface markers.

The antibody staining concentrations were determined by titration on positive and negative control cell populations. Specifically, intra-nuclear antibody concentrations were calibrated using MCF 7 cells line (*TP53* WT) and RXF 393 cell line (*TP53*^R175H^). Absence of false positive staining was validated using HL-60 cell line (*TP53* null).

Intra-cellular antibodies that were purchased from different batches were combined, and the titer of the combined pool was validated (with the first, calibrated batch, serving as a reference) using a primary AML sample harboring *TP53*^P151A^ mutation. Since cadmium and zinc chelators can influence the conformation of p53^4^, all the reagents that were used for the intra-cellular staining did not contain zinc chelators, and antibodies were not conjugated to cadmium.

All samples were stained and recorded in duplicates (except for the mobilized PBMCs donations, due to paucity of available cells).

Xenotransplantation assays and *in vivo* pharmacologic treatment

We used the following mouse strains of immune-deficient NSG (NOD/SCID/IL-2Rgc-null) mice: NSG (Stock No: 005557), NSG-hSCF (Stock No: 017830), that transgenically express human SCF, and NSG-SGM3 (Stock No: 013062), that transgenically express human IL-3, GM-CSF and SCF (all from The Jackson Laboratory, Bar-Harbor, ME, USA).

1-2.5x10^6^ CD3 depleted mononuclear cells were injected intra-femoraly (right femur) into 8 to 12-week-old female mice. Mice were sub-lethally irradiated (200 cGy) 6–24 hours before human cells’ injection.

pCAP-250 and the control peptide were delivered by implanting subcutaneous micro-osmotic pumps that excrete the peptides at a rate of 19 mg/kg/day (over a period of 14 days), achieving blood concentrations of approximately 6 ugr/mL^5^.

Mice were sacrificed on day 56 (three weeks following pump implantation). The bone marrows of the injected bone (right femur) and the non-injected bones (left femur, tibiae) were flushed with Iscove's Modified Dulbecco's Medium (IMDM) (Cat: 01-058-1A, Biological Industries, Beit Ha’emek, Israel). Cells were filtered through a 35μm cell strainer (Cat: 352235, Corning, Corning, NY, USA) to obtain a single-cell suspension.

Human engraftment was assessed by flow cytometry, as described below.

Experiments that included RNA sequencing and mass cytometry analyses of engrafting cells were performed as follows: eight weeks following human AML sample injection, mice were IV injected with a single dose of either pCAP-250 or the scrambled peptide (16.6 mg/kg). Mice were sacrificed 12 hours later. Bone marrows were harvested as described.

Flow Cytometry

Analyses were performed using antibody panel (supplemental table 3), on Cytoflex (Beckman Coulter, Brea, CA, USA), using CytExpert software v 2.4.0.28 (Beckman Coulter, Brea, CA, USA).

Deep targeted DNA Sequencing

Sequencing of mononuclear hematopoietic cells and of expanded T cells from each sample was performed in duplicates following genomic DNA extraction (DNeasy kit, Qiagen, Hilden, Germany).

We used a panel of single molecule Molecular Inversion Probes (smMIPs)^6^ (designed with MIPgen software^7^ that covers recurrently mutated AML “hotspots” in 33 genes (supplemental table 4).

For deep targeted DNA sequencing of engrafting human cells, cells retrieved from bone marrows of sacrificed mice were sorted by FACS BD FACSAria™ III Cell Sorter (BD Biosciences, San Jose, CA, USA) according to the main engrafting sub-populations using a panel of monoclonal antibodies (supplemental table 5). Sorted cells underwent whole genome amplification (Repli-G, Qiagen, Hilden, Germany).

Libraries were prepared using a panel of smMIPs that was designed to capture mutations known to be present in the original injected samples (supplemental table 6).

Validation of sequencing results of *DNMT3A*^R882^ was performed when the coverage depth of the targets was insufficient (less than 100X). This was done by preparing a different library using an amplicon-based approach. This approach was used also when only a single target was of interest (i.e.: for *DNMT3A*^R882^ mutated clonal hematopoiesis samples).
Primary PCR for *DNMT3A* (exon 23) was performed with the following primers:
Forward:CTACACGACGCTCTTCCGATCTTAACTTTGTGTCGCTACCTC
Reverse:CAGACGTGTGCTCTTCCGATCTTTTTCTCCCCCAGGGTATTTG
Secondary PCR was performed with the following primers:

Forward primer:

AATGATACGGCGACCACCGAGATCTACAC[Fw_Index_D5XX]ACACTCTTTCCCTACACGACGCTCTTCCG;

Reverse primer:

CAAGCAGAAGACGGCATACGAGAT[Rev_Index_D7XX]GTGACTGGAGTTCAGACGTGTGCTCTTCCG;

All Sequencing were performed with MiSeq, MiniSeq and NovaSeq sequencers (Illumina, San Diego, CA, USA).

Data pre-processing and variant calling

Paired-end 2 X 151bp sequencing data were converted to fastq format. Reads were merged using BBmerge v38.62^8^ with default parameters, followed by trimming of the ligation and extension arm using Cutadapt v2.10^9^. Unique Molecular Identifiers (UMIs) were trimmed and assigned to each read header. Processed reads were aligned using BWA-MEM^10^ to a custom reference genome, comprised of the appropriate smMIP panel sequences ± 150 bases extracted from broad hg19. Aligned files were sorted, converted to BAM (SAMTools V1.9^11^) followed by Indel realignment using AddOrReplaceReadGroups (Picard tools) and later IndelRealigner (GATK v.3.7^12^). Variant calling was done using mpileup and MuTect2 (in a tumor-only mode, GATK 3.7^12^ (for the single nucleotide variant (SNVs), and Varscan2 v2.3.9^13^ and Platypus^14^ for indels. Variants were annotated using ANNOVAR^15^. This bioinformatics pipeline can identify variants as low as 0.005 (manuscript under preparation).

Single cell RNA sequencing (scRNAseq)

Libraries were prepared using 10X Genomics Chromium Single Cell 3ʹ Reagent Kits v3 (10X Genomics, Pleasanton, CA, USA) according to manufacturer’s protocol. Libraries were prepared twice, using biological duplicates. Sequencing was performed with Illumina NextSeq sequencer (Illumina, San Diego, CA, USA).

Demultiplexing, alignment, filtering, barcode counting, and UMI counting were performed using Cell-Ranger (version 3.1.0) bioinformatics pipeline.

In order to exclude contamination of mice cells in the samples that were obtained following engraftment, the pipeline was run twice with different genomes: once with a joint genome of mm10 and hg38 and the other with only hg38. Cells were determined to be of murine origin if the expression of mouse genes was above 20% from the total gene expression in the joint analysis (mm10 & hg38).

Altogether, 80-120 cells were excluded from each sample.

The output from the hg38 genome was used for further analysis with R version 3.6.0 and Seurat version 3.1.1^16^.

Cell filtering was based on the total number of genes or UMI counts per cell (high or low 5 percentiles were removed), and the percentage of mitochondria genes (filtered out when higher than 20%). The number of cells after filtering ranged between 2203 and 3779 per sample. 26 clusters were created with 2000 variable genes and 20 principal components (PCs).

Additionally, Somatic variants were called for genes that are known to be mutated in the samples.

Gene set enrichment analyses

An overrepresentation analysis (ORA)^17^ was employed. The gene sets associated with a blood or immune cell types, were downloaded from PanglaoDB database^18^ and compared to the differentially positive expressed cluster markers filtered with criteria of lnFC ≥ 1 and padj ≤ 0.05. To test for overrepresentation of successes in the sample, the hypergeometric p value was calculated using R function phyper with lower tail= false as the probability of randomly drawing k or more successes from the population in n total draws^19^. The FDR was achieved by adjusting the p value using Benjamini and Hochberg^20^. In addition, Enrichr was used to identify enrichment of cluster marker genes with various molecular pathways^21,22^.

Statistical analyses

Dependency between treatment and the relative size of each cluster was determined by chi square test for independence. For a two-by-two table the Yates continuity correction was applied. A post-hoc standardized residuals analysis with Bonferroni correction for multiple comparisons was used to find out which clusters had a significantly different size than expected.

**Supplemental figures**

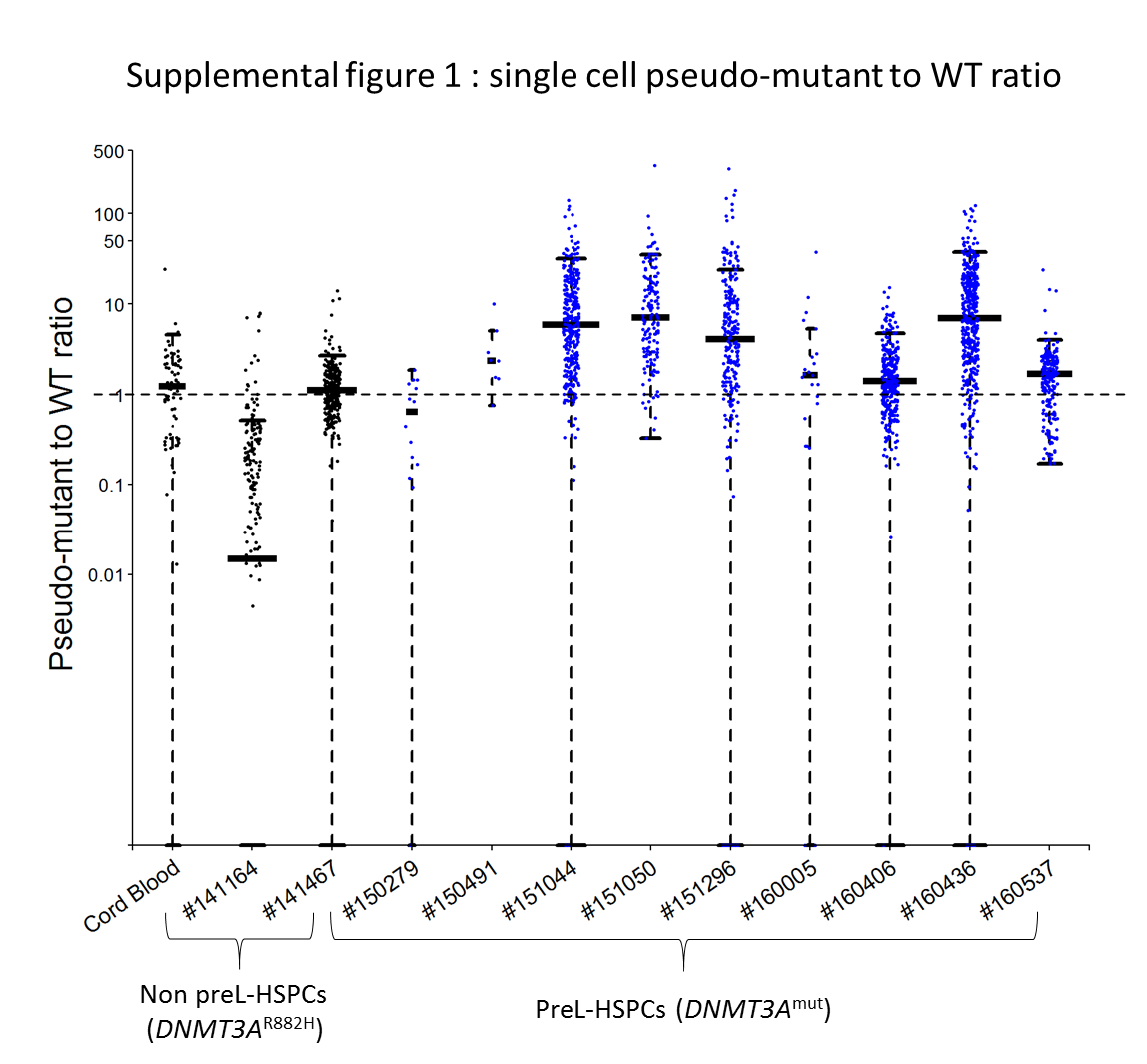

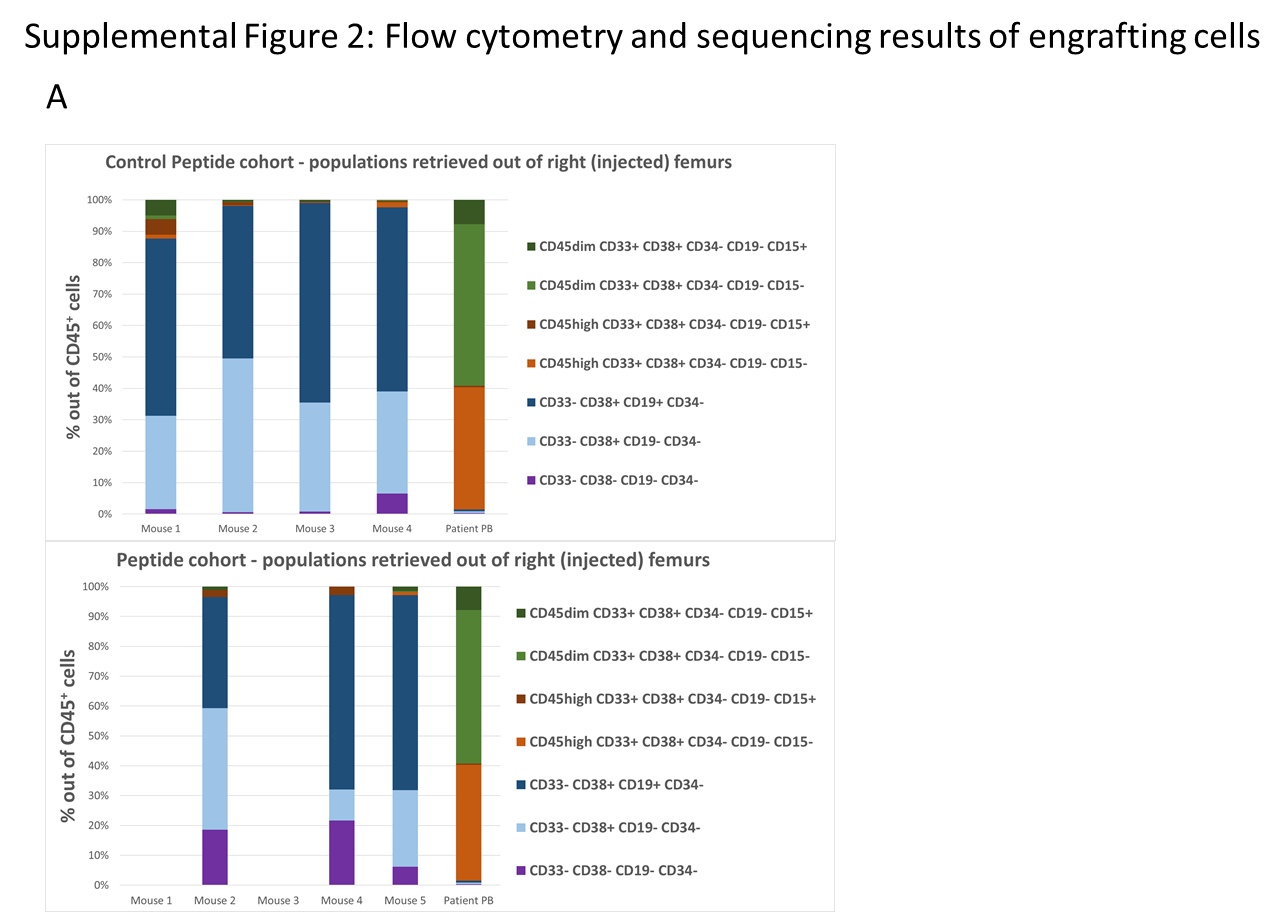

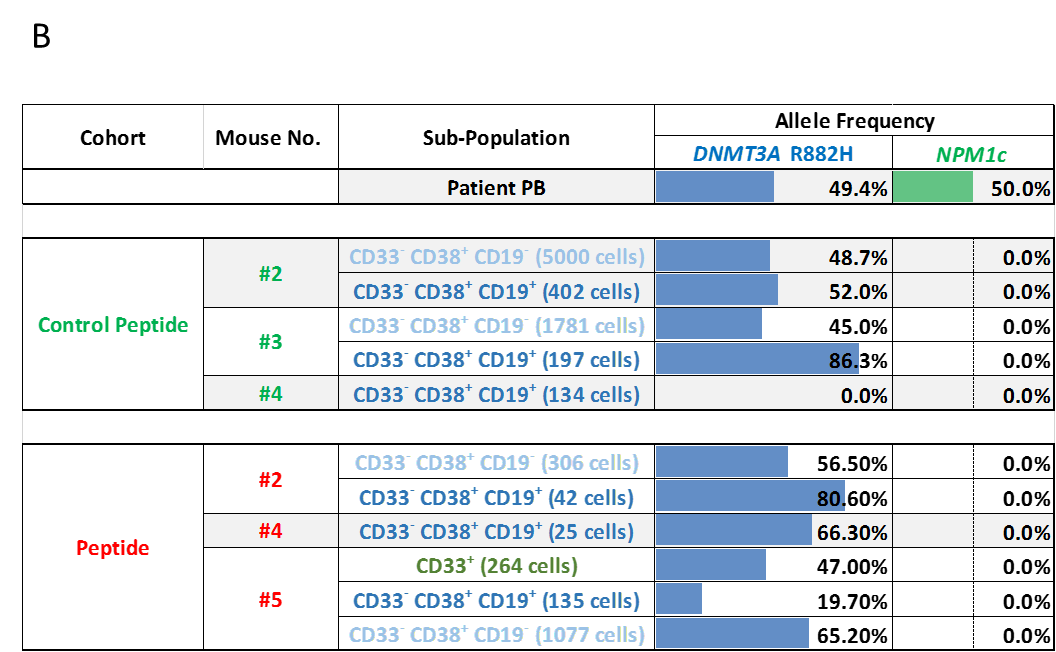

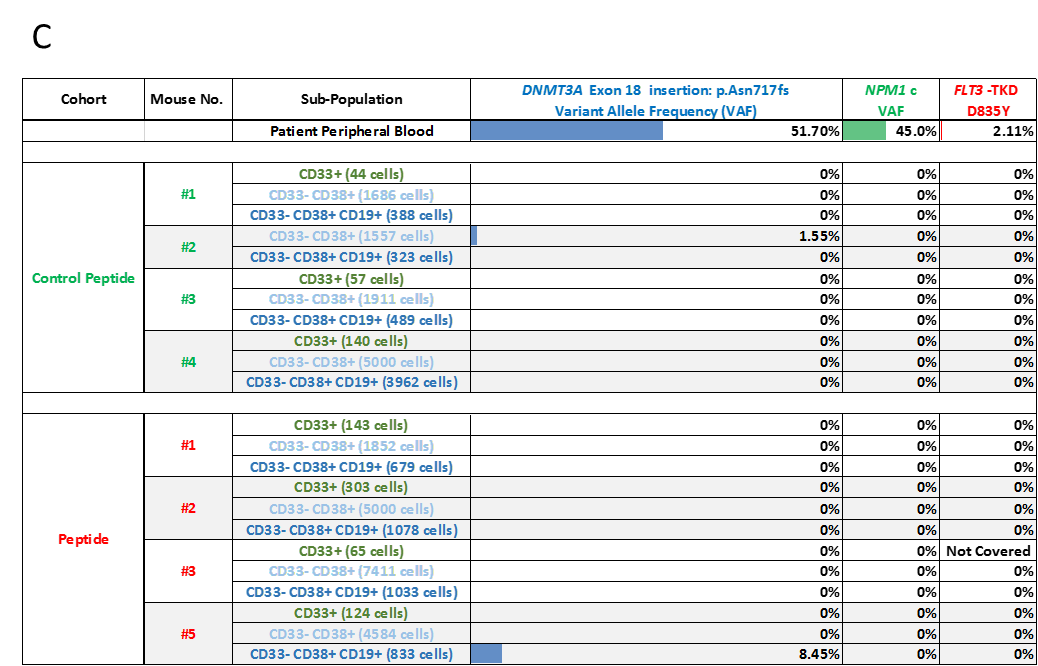

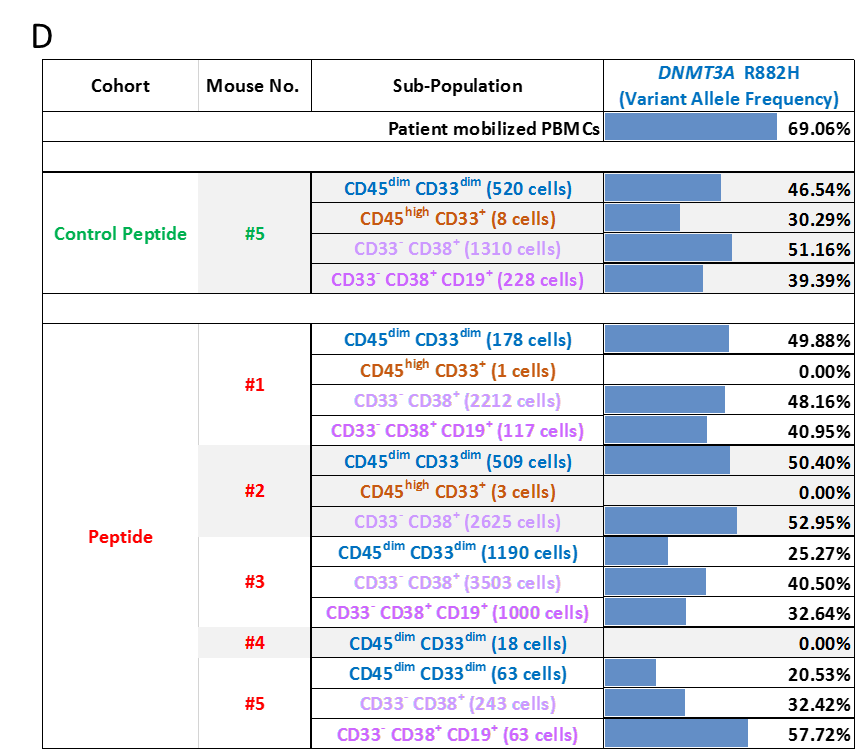

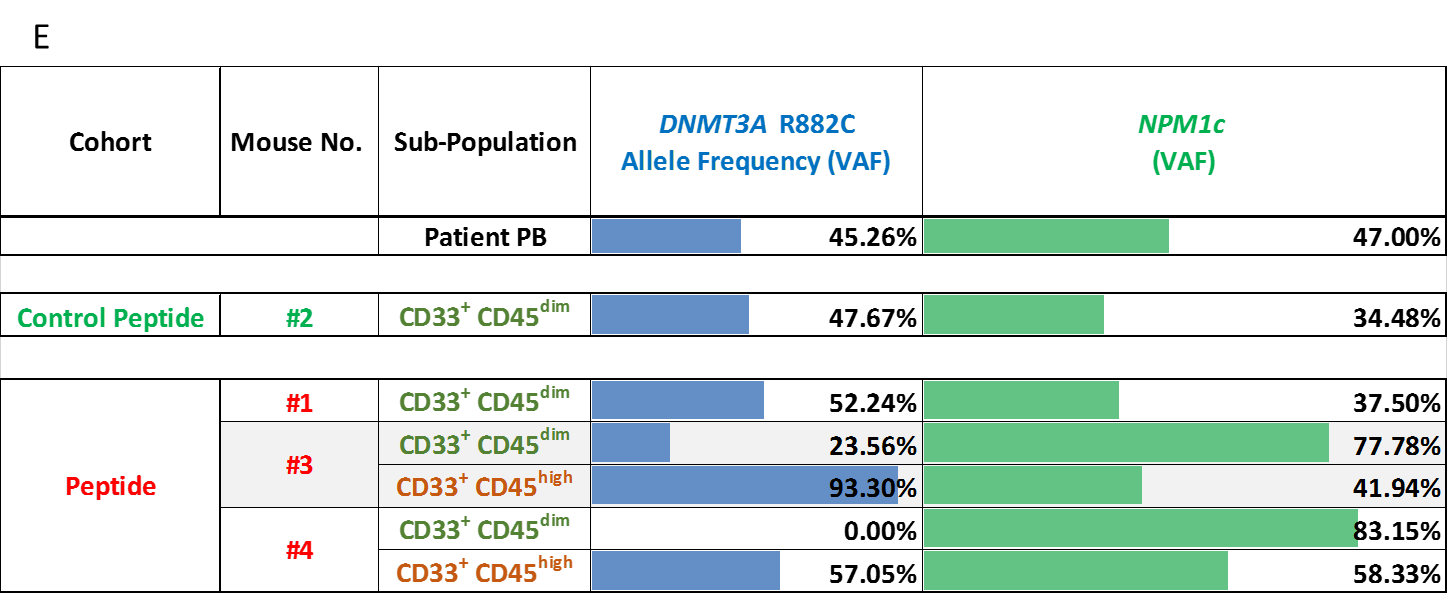

**Legends for supplemental figures**

Supplemental Figure 1:

Ratio between the pseudo-mutant conformation and the WT conformation of p53 at the single cell level in different samples.

From left to right: HSPCs from a cord blood sample, HSPCs from 2 samples of *TP53* WT, *DNMT3A*^R882H^ CH (all in black). HSPCs from 9 *TP53* WT, *DNMT3A*-mutated, AML samples (blue).

In eight out of nine preL-HSPCs samples the dominant conformation of p53 is the pseudo-mutant. This ratio was around or below 1 (dashed line) in HSPCs from non-leukemic samples.

The median and 1.5 X interquartile range are presented for each sample.

The medians are drawn proportional to the square-roots of the number of observations in each cohort.

Supplemental Figure 2:

(A) Flow cytometry analyses of representative engrafting sub-populations of sample #160005 (Figure 2B). Most engrafting cells are non-myeloid (blue colors) as opposed to the injected blasts (right columns), reflecting their differentiation capacity and their non-leukemic origin.

(B) Sequencing results of the engrafting sub-populations of *DNMT3A*^R882H^, *NPM1*c AML (sample #160005, Figure 2B) that are presented in (A).

The number of sorted cells in each sub-population is presented.

The patient’s (injected) cells are presented in the first row. Except for a single sub-population, all engrafting cells are pre-leukemic harboring only *DNMT3A*^R882H^ mutation, without *NPM1*c mutation.

(C) Sequencing results of engrafting sub-populations of *DNMT3A*mut, *NPM1*c, *FLT3*-TKD AML sample (Figure 2D).

The number of sorted cells in each sub-population is presented.

This sample gave rise to a multi-lineage graft. Except for a few, rare cells, most engrafting cells do not harbor any of the mutations that were found in the injected, leukemic cells (first row). Therefore, engrafting cells originated from a non-pre-leukemic clone (or clones).

(D) Sequencing results of engrafting sub-populations of a *DNMT3A*^R882H^ CH (Figure 2E).

The number of sorted cells in each sub-population is presented.

Except for a few sub-populations, all engrafting cells harbor the *DNMT3A*^R882H^ mutation.

(E) Sequencing results of engrafting sub-populations of a *DNMT3A*^R882C^, *NPM1*c, *FLT3*-ITD (low allelic ratio) AML sample. 5000 cells were sorted from each sub-population.
Engrafting cells originated from leukemic stem cells since they all harbor *NPM1*c mutations, similar to the injected cells (first row).

PB – peripheral blood, PBMCs – peripheral blood mononuclear cells.

**Supplemental tables**

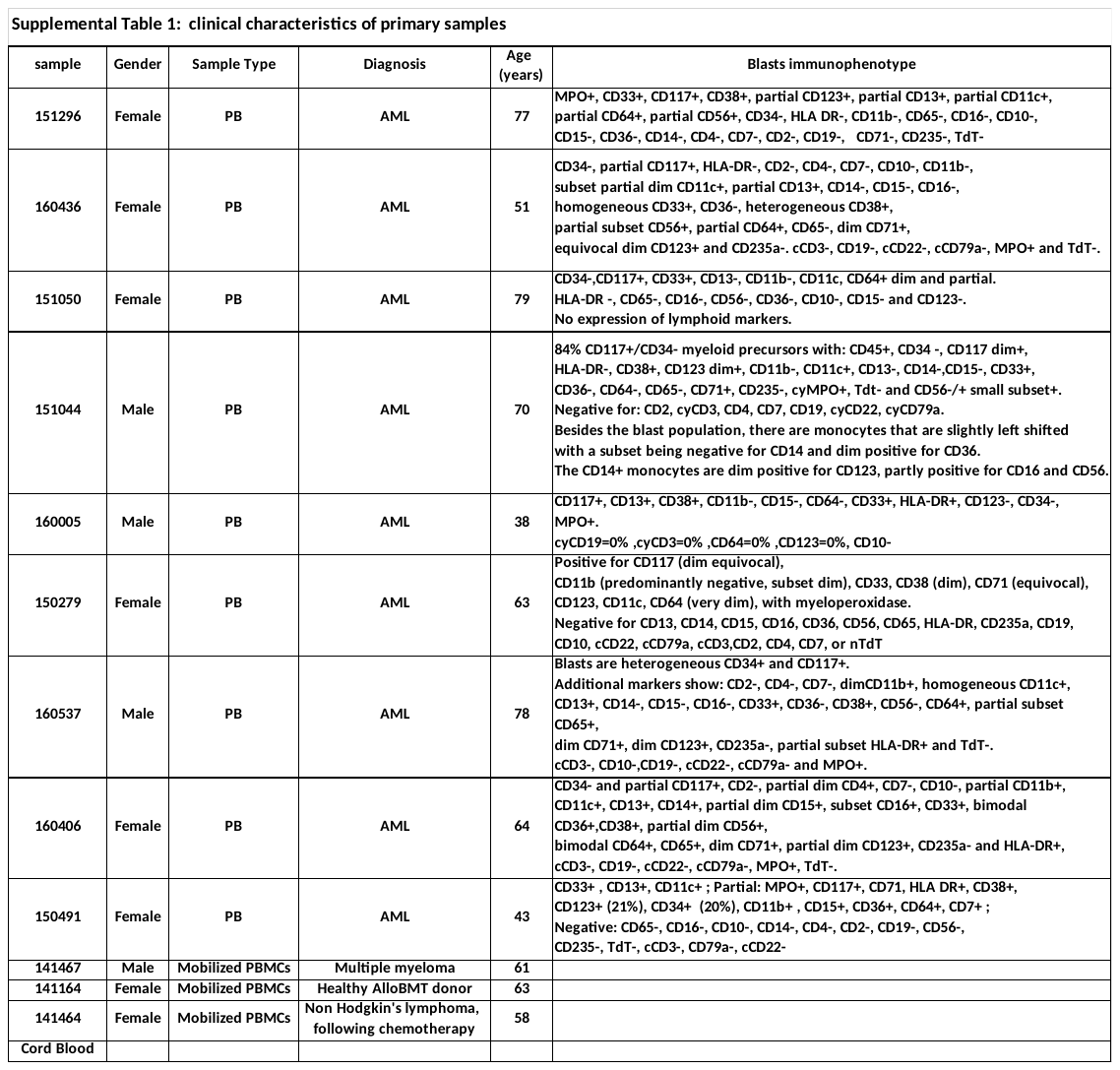

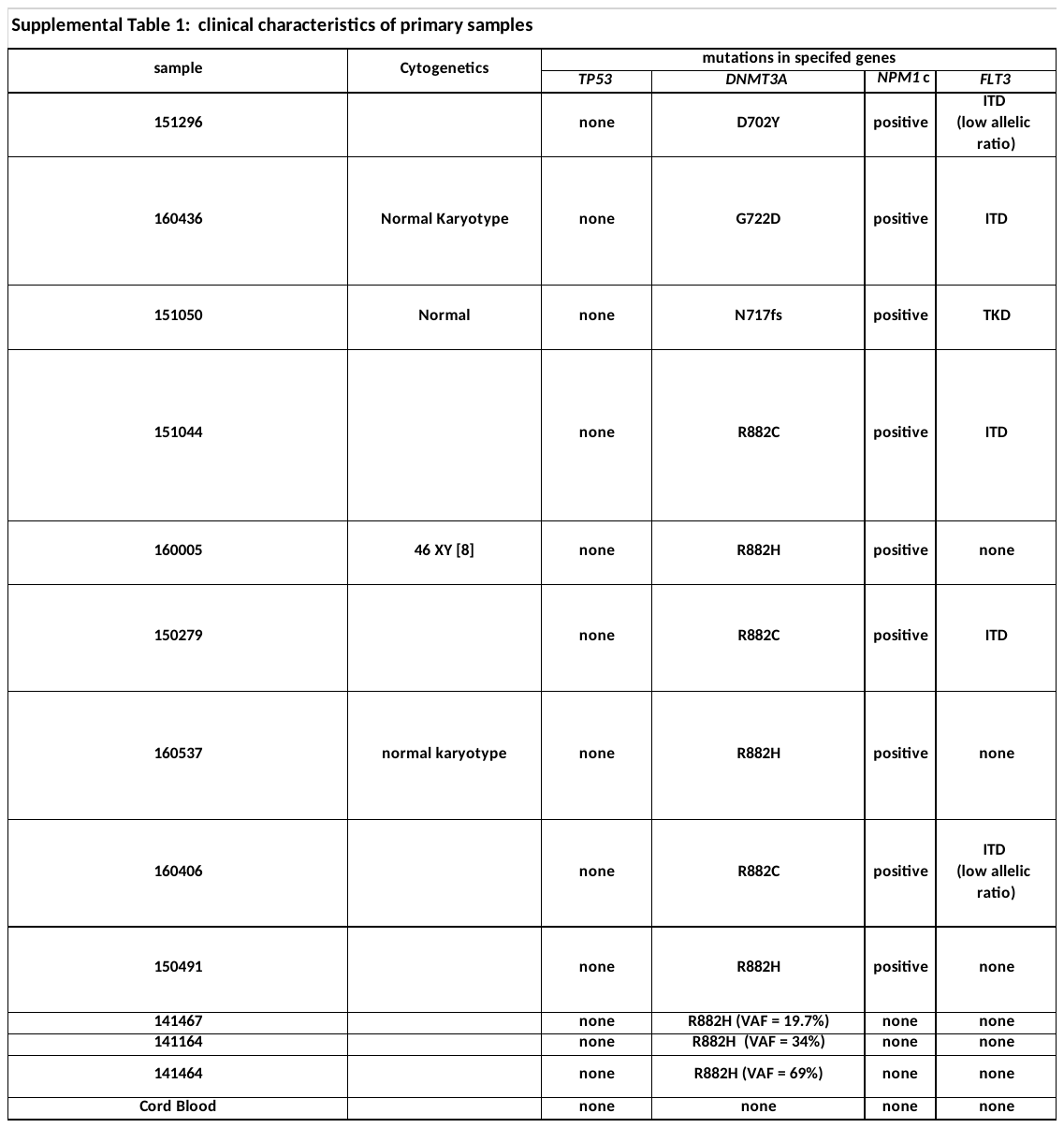

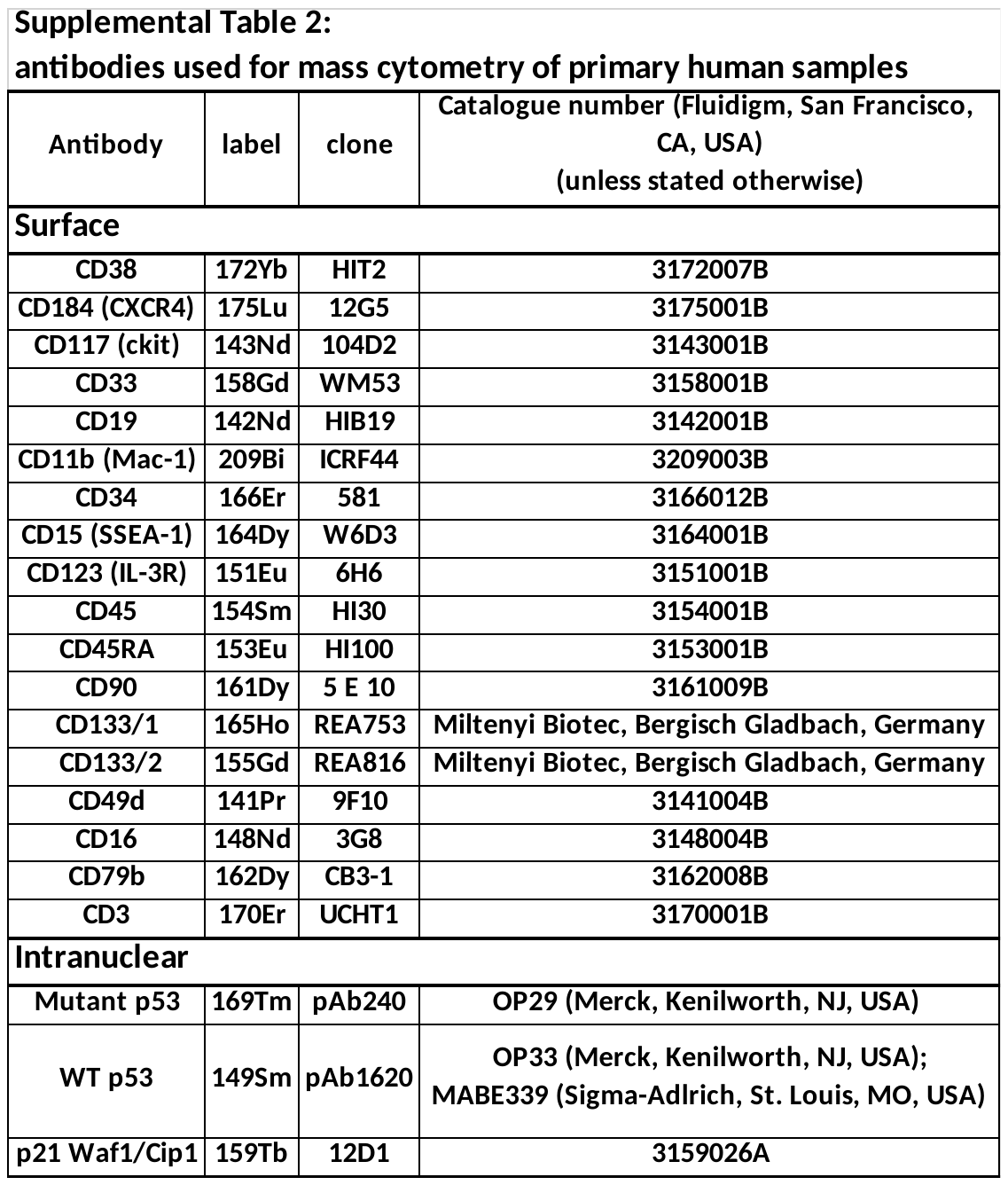

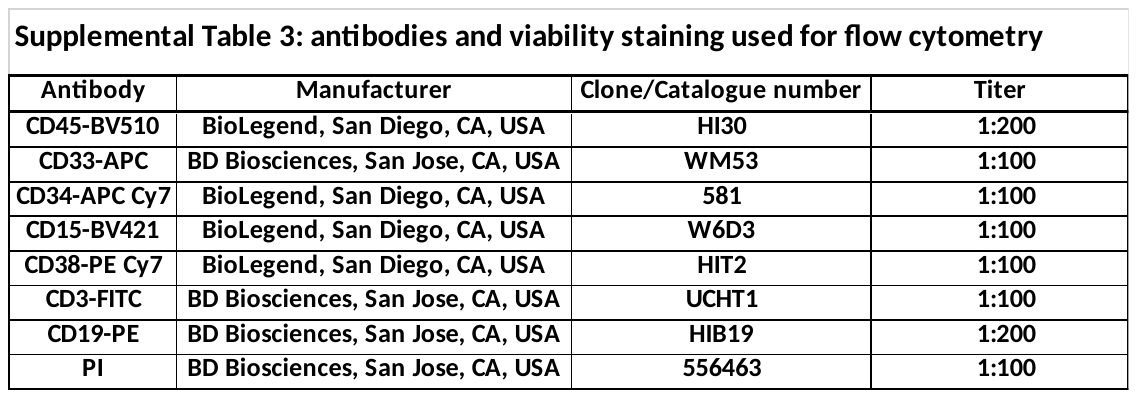

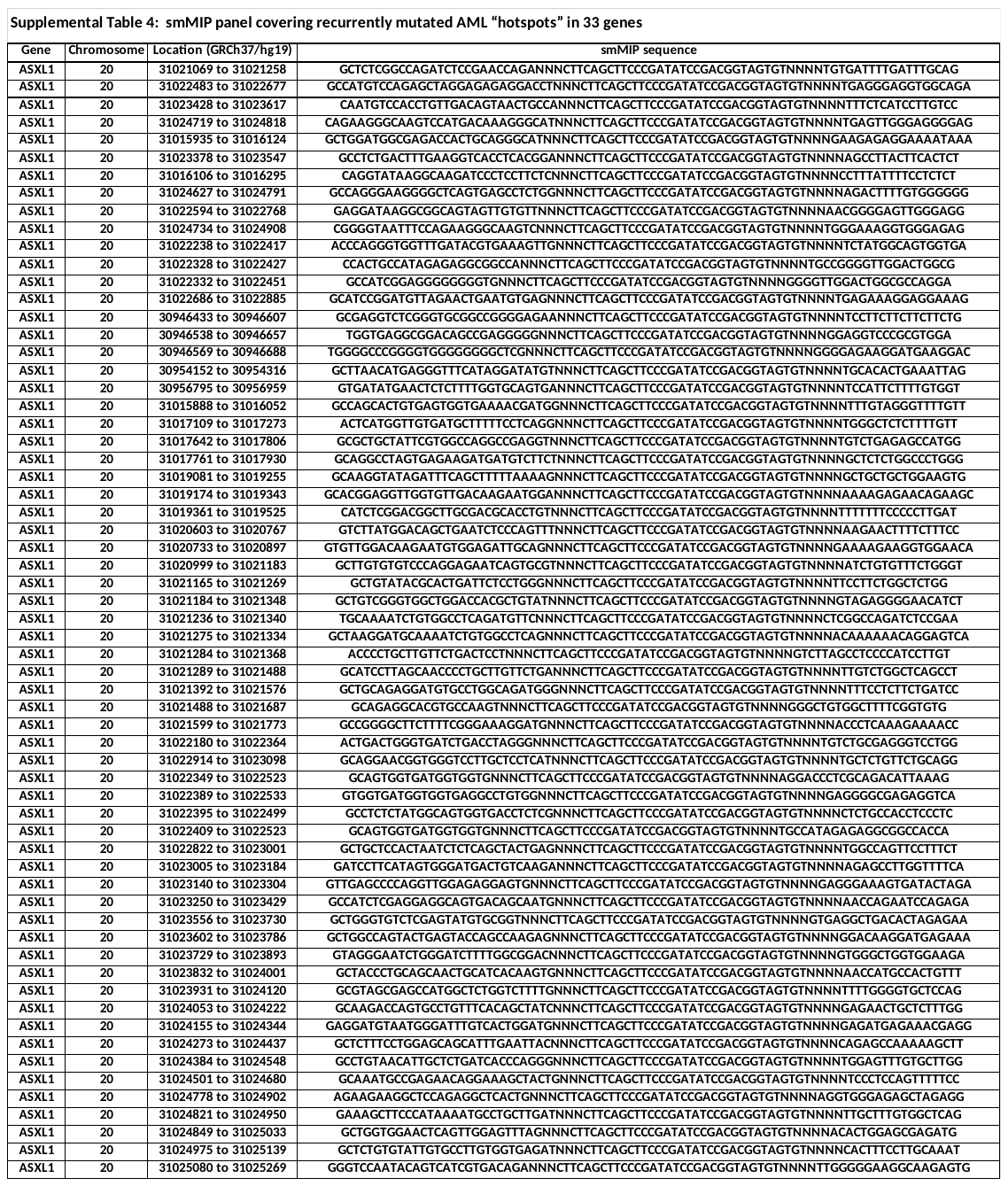

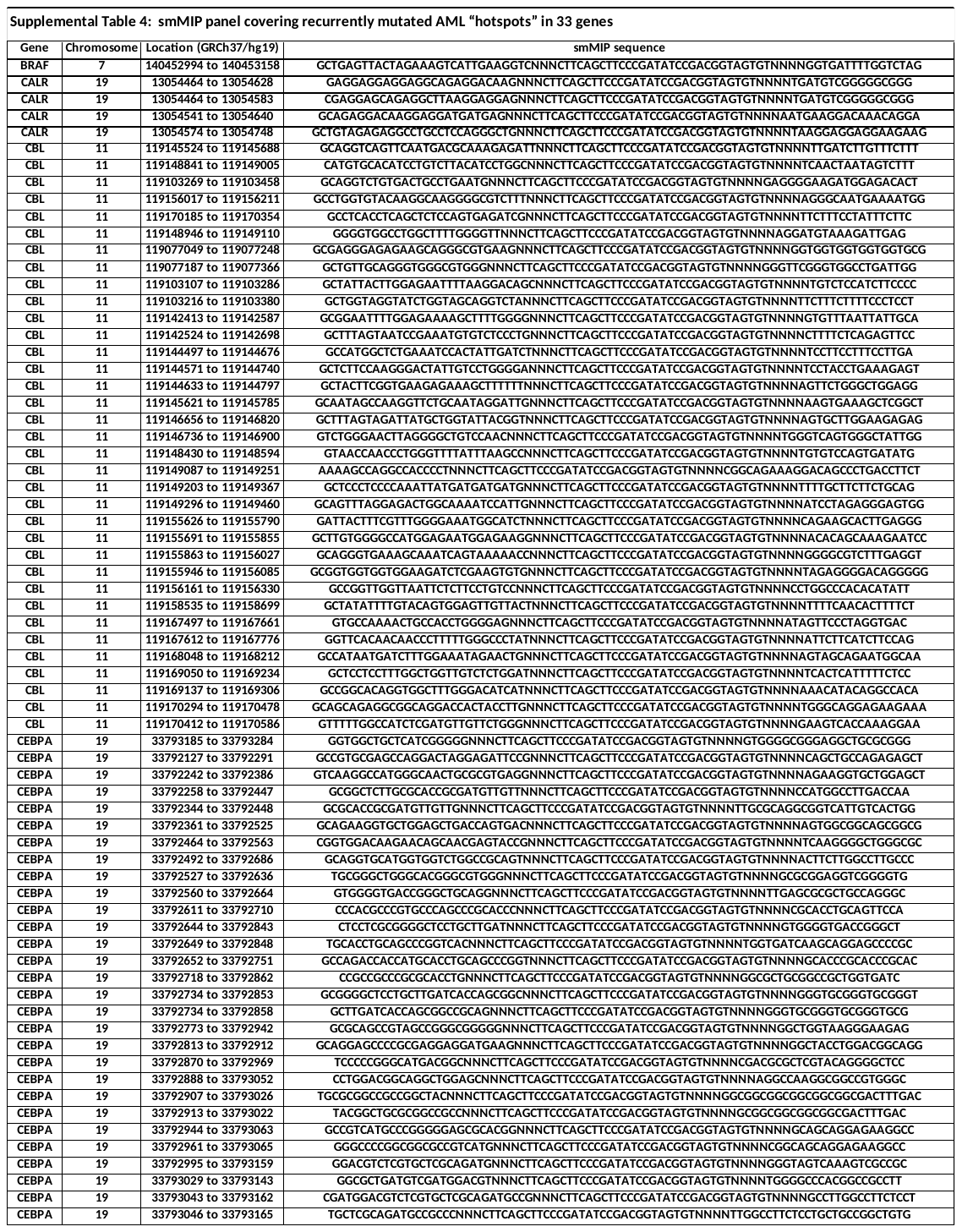

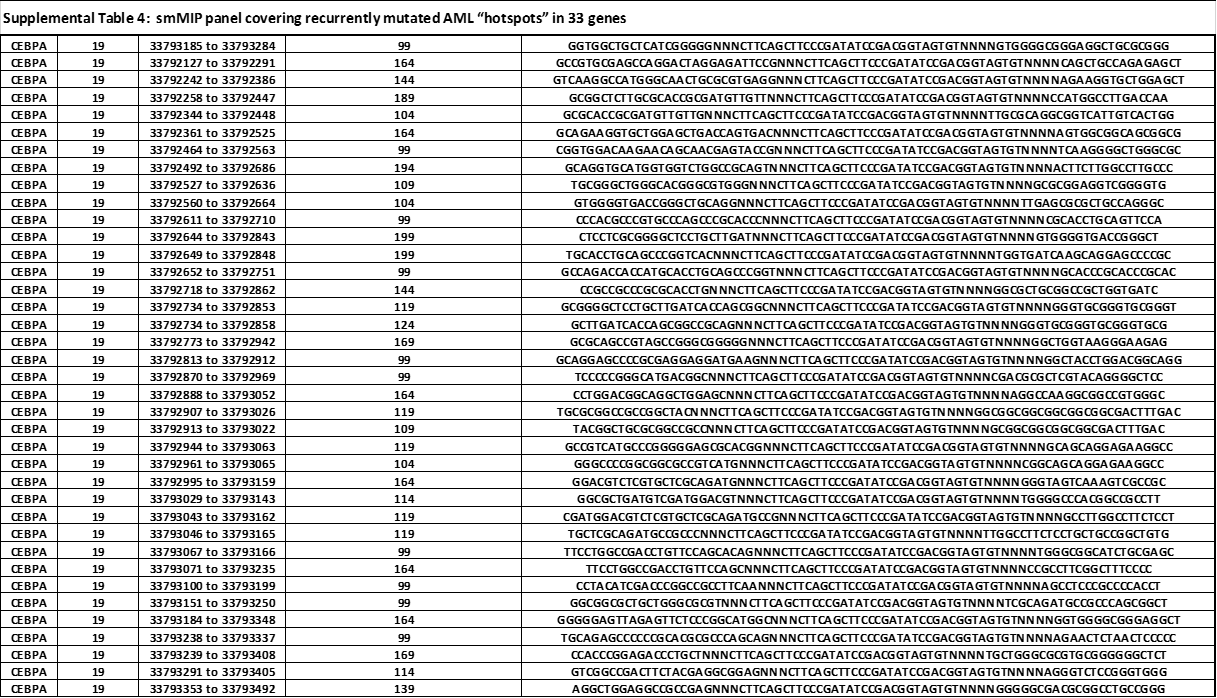

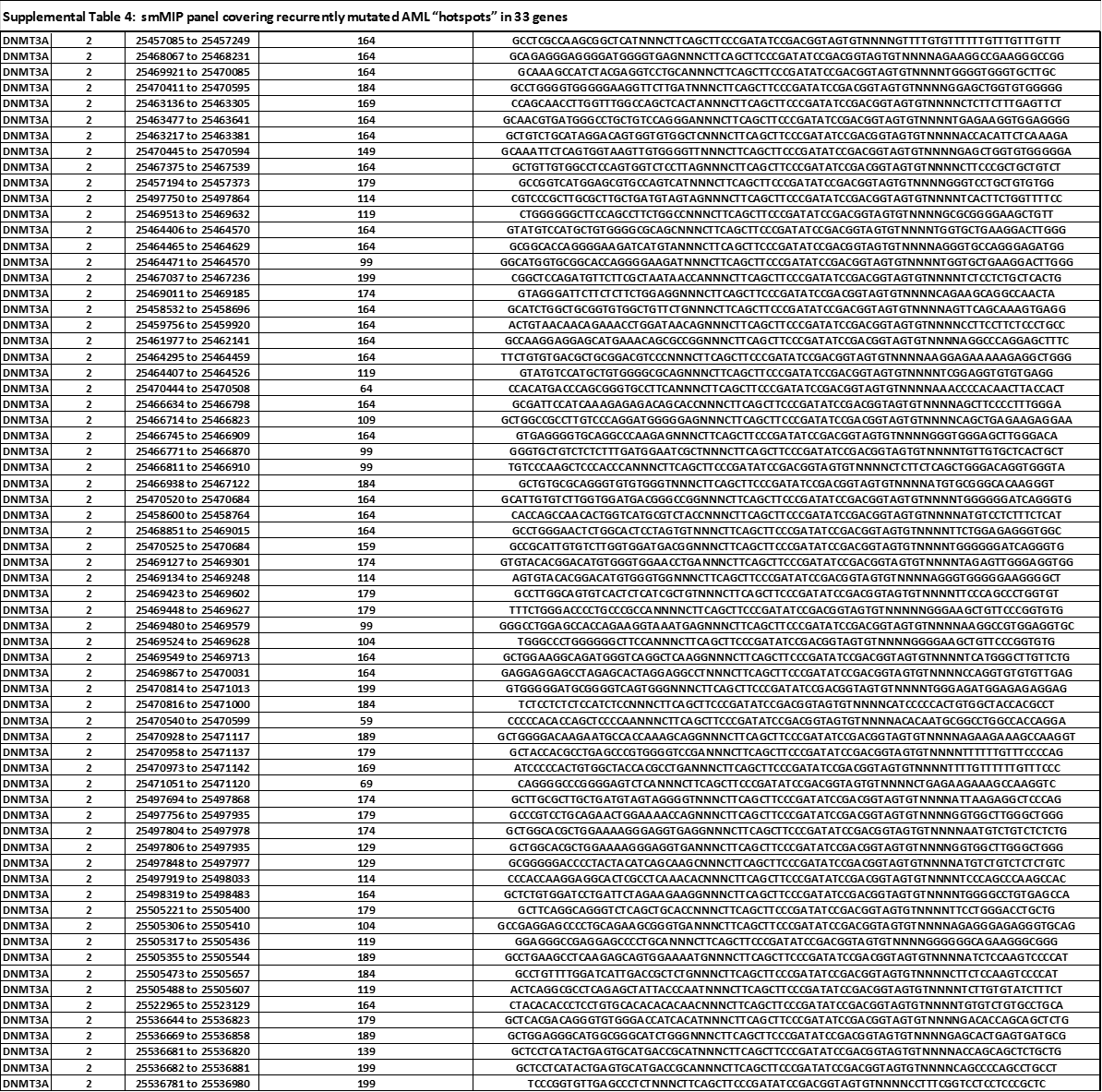

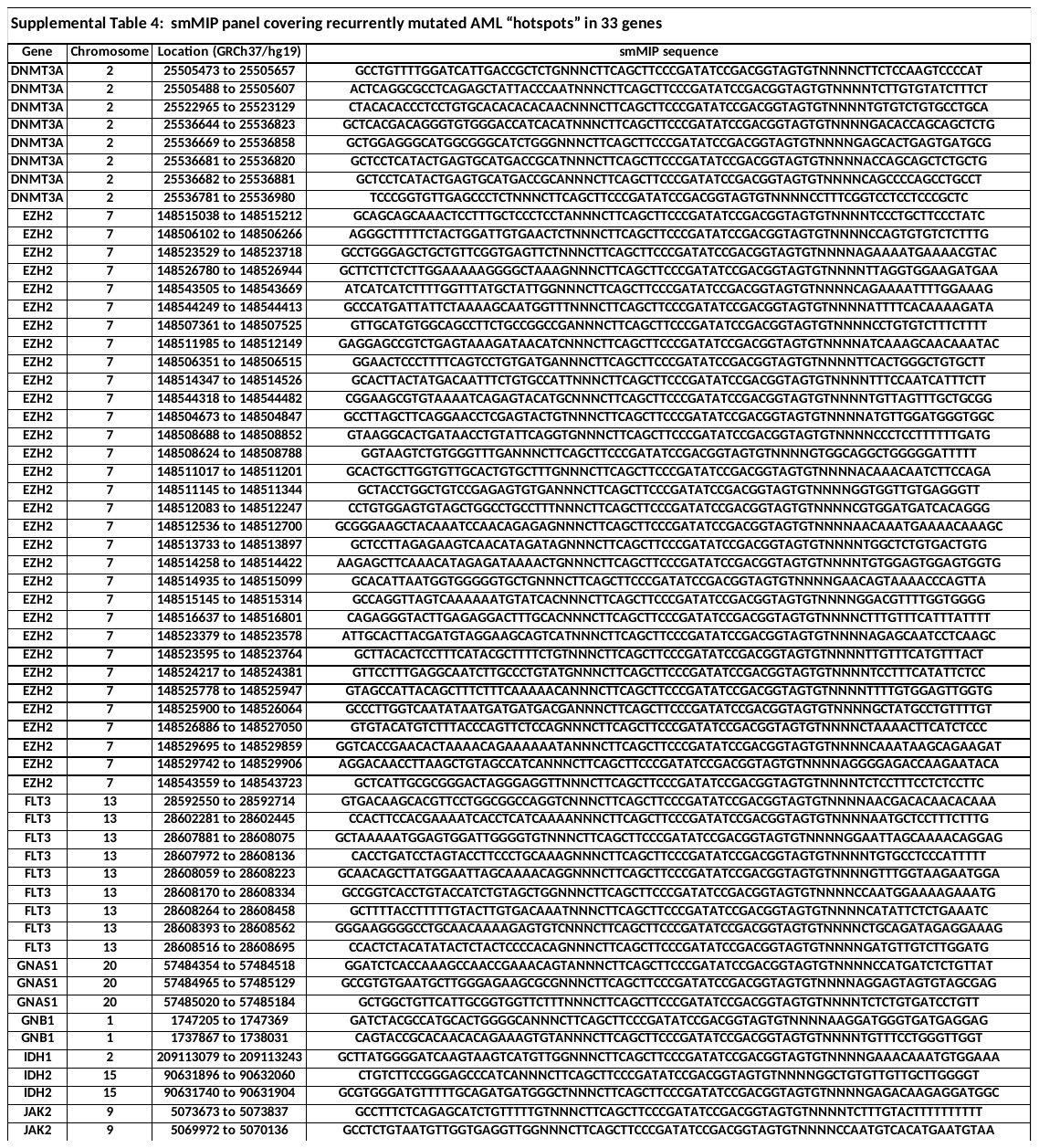

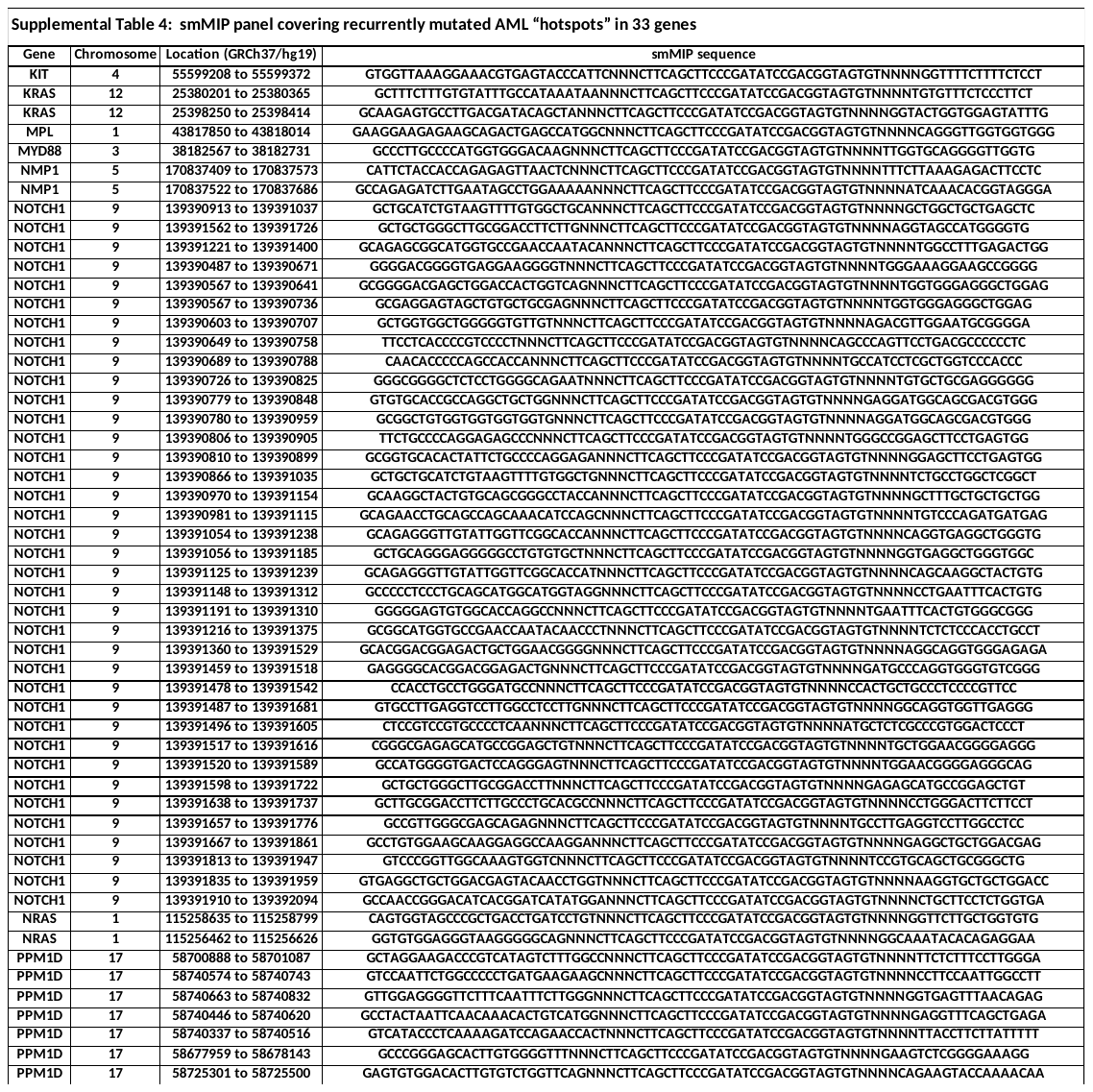

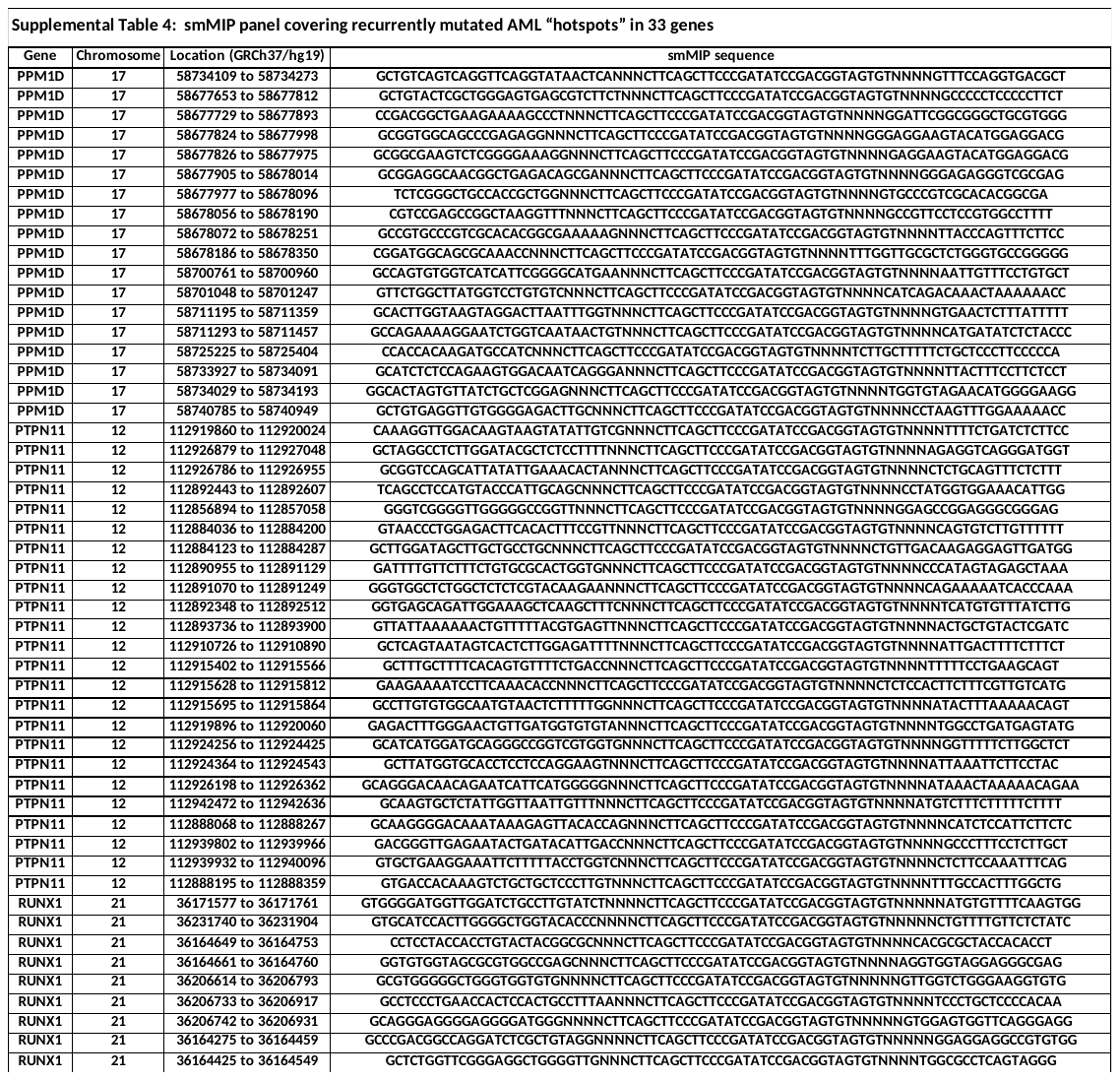

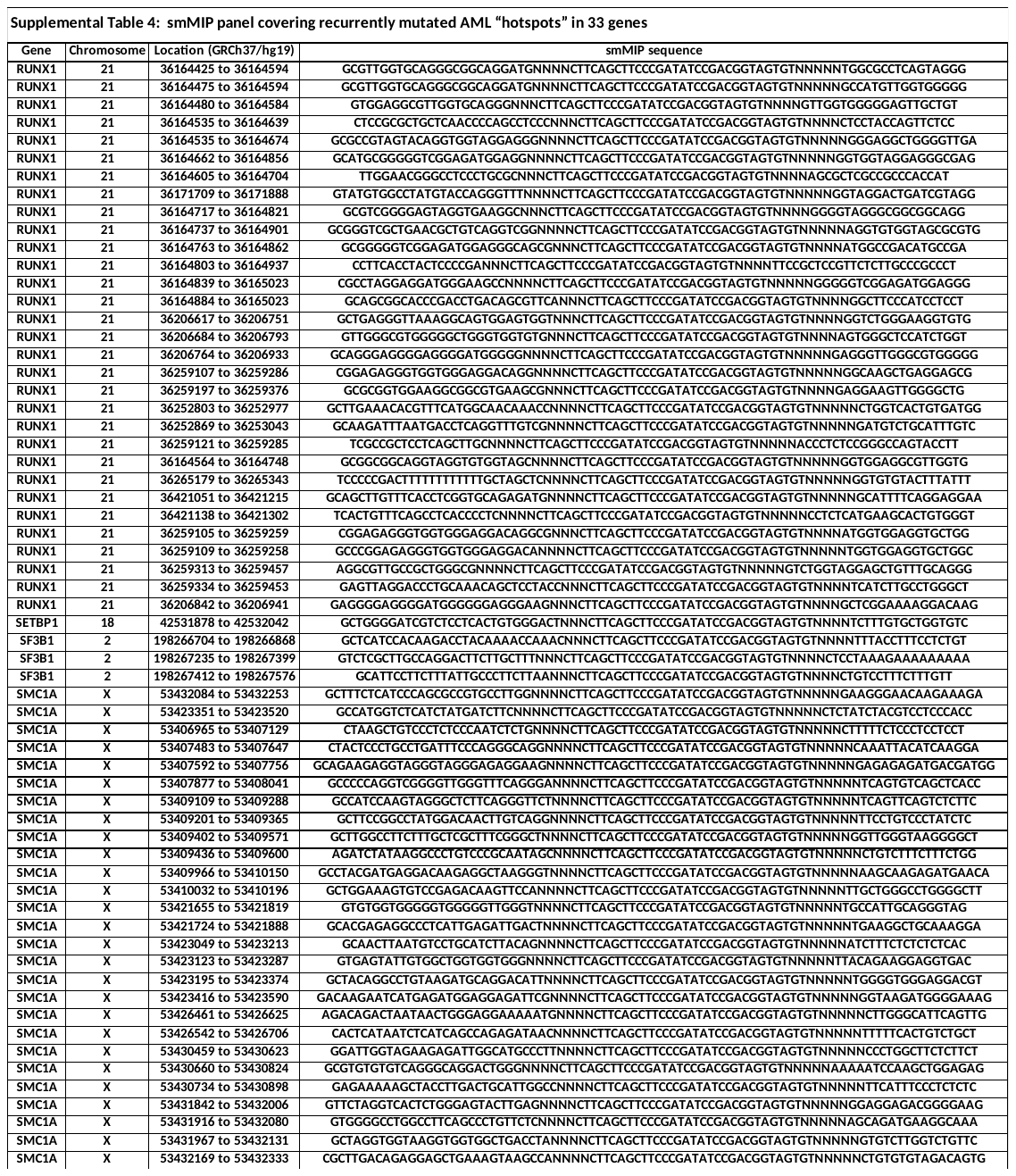

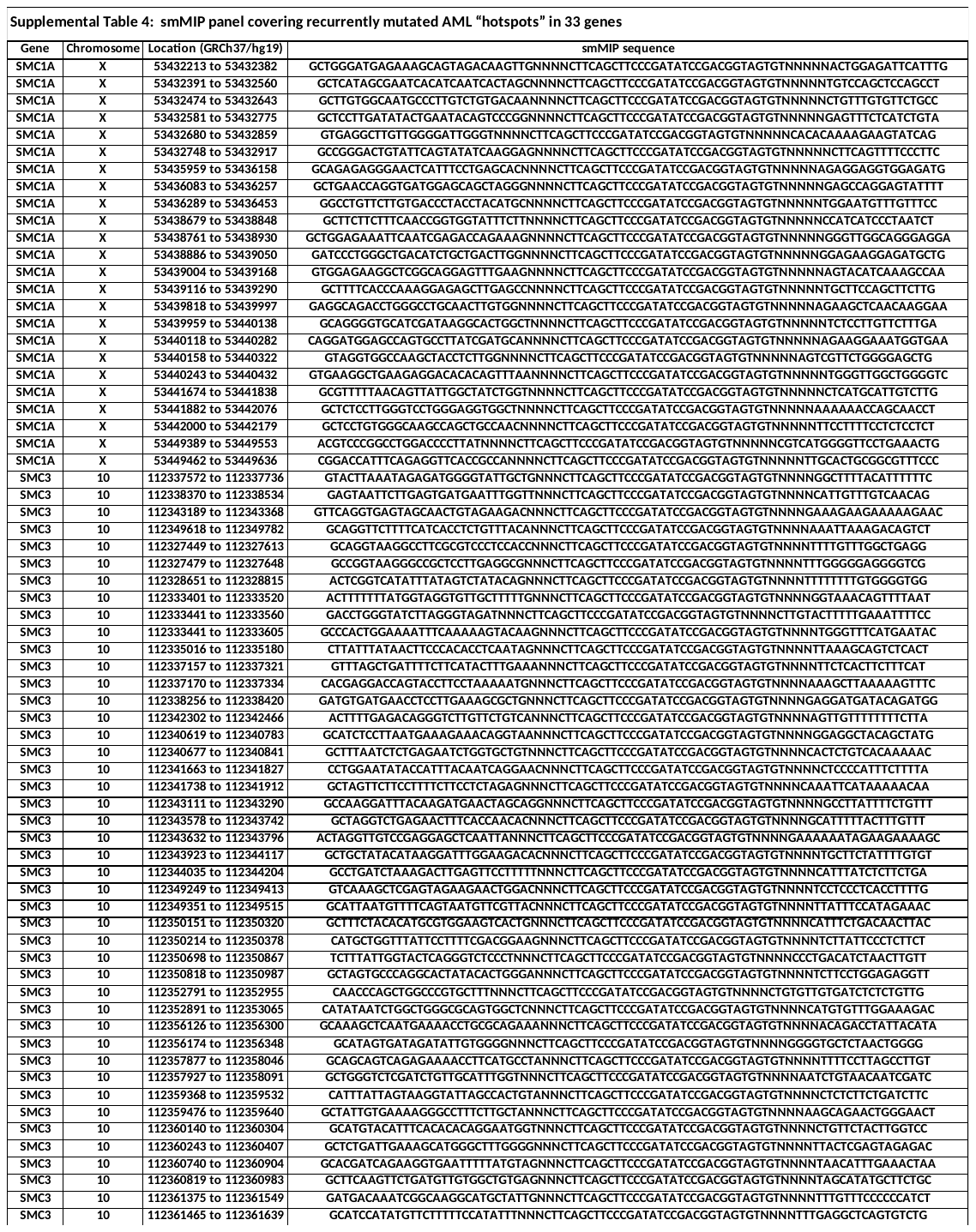

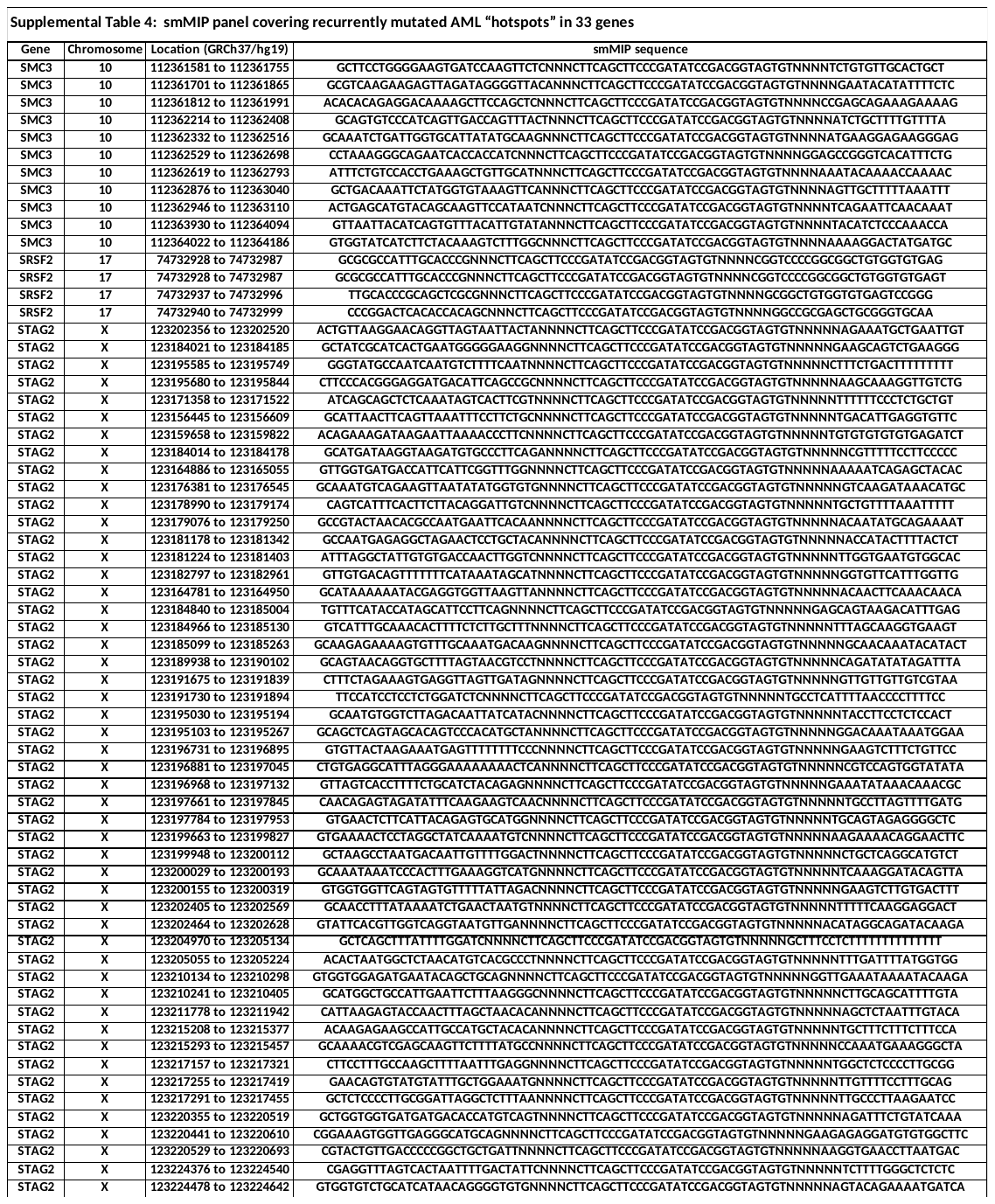

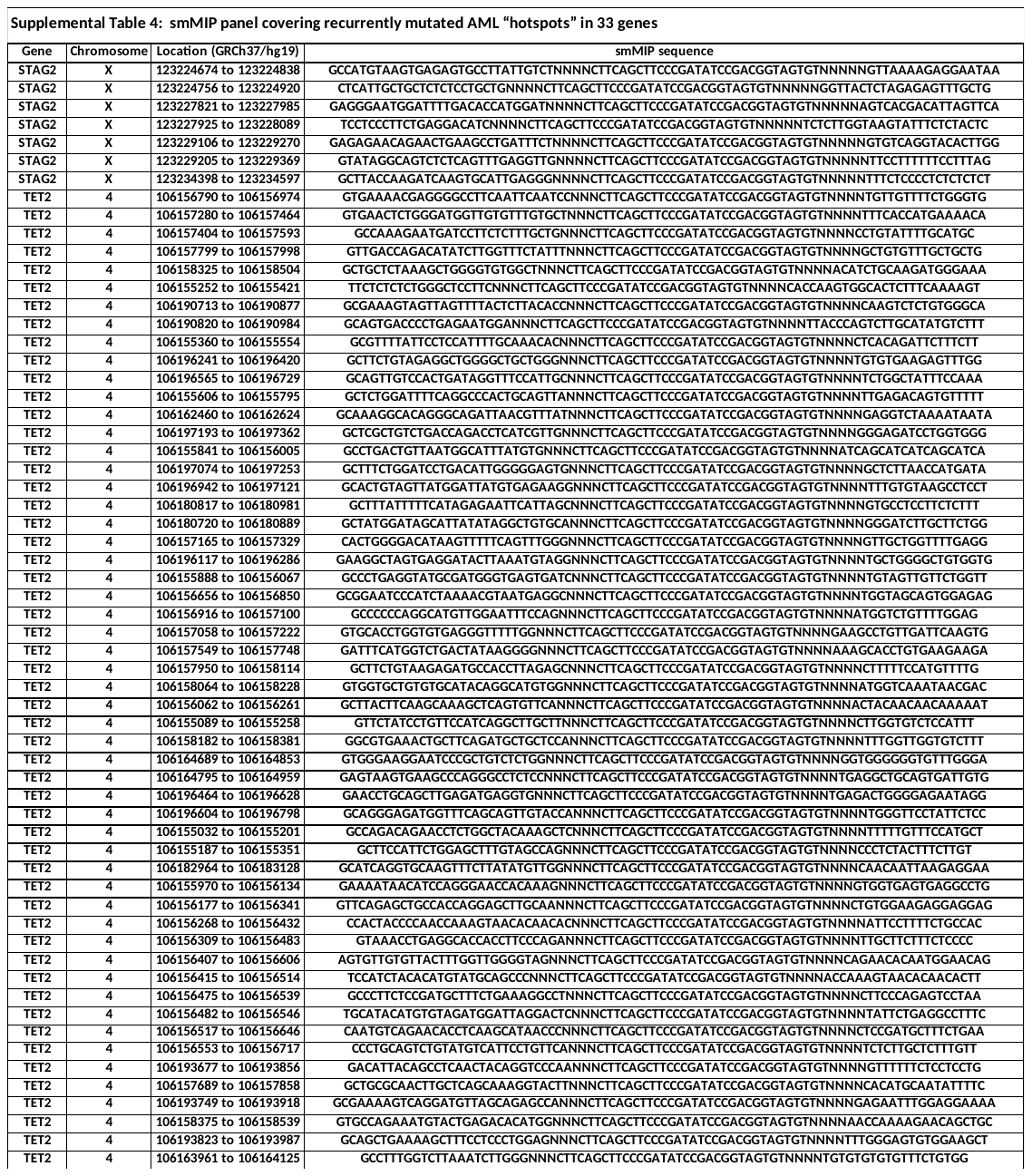

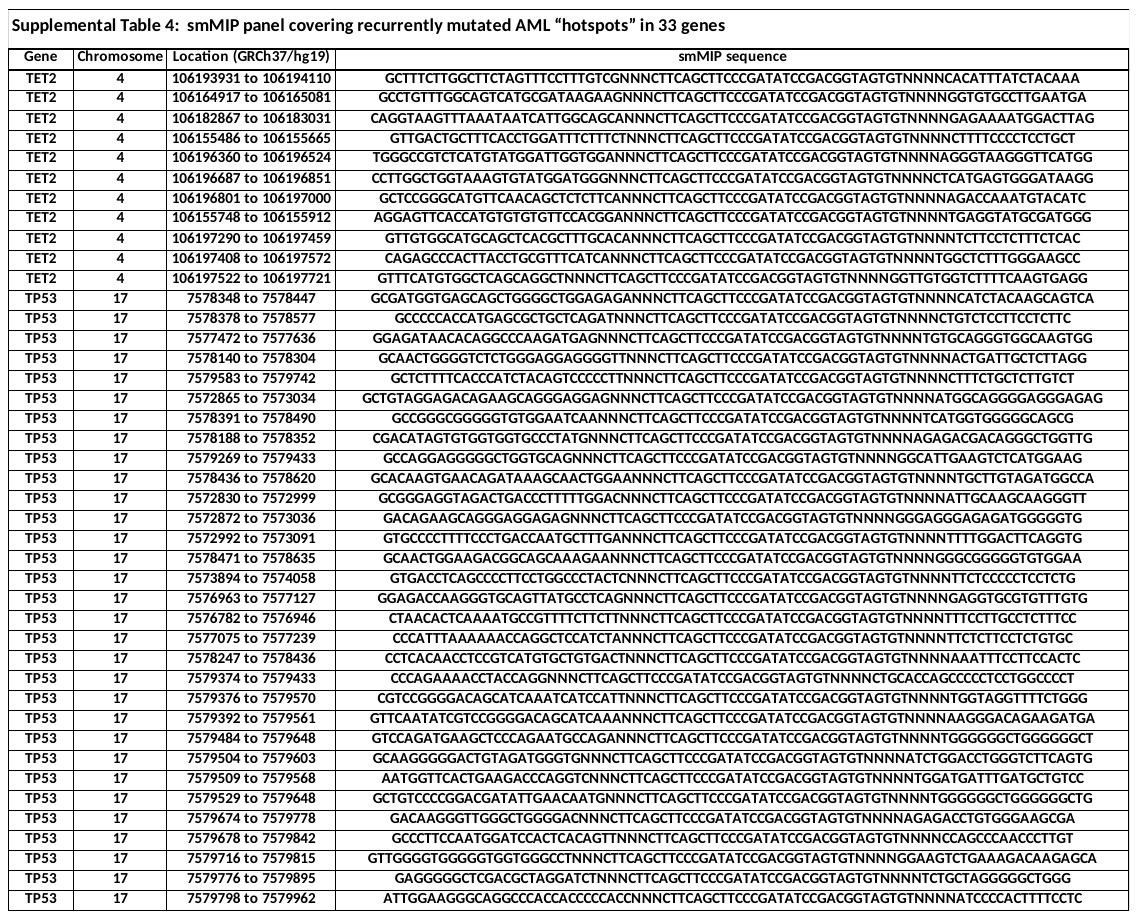

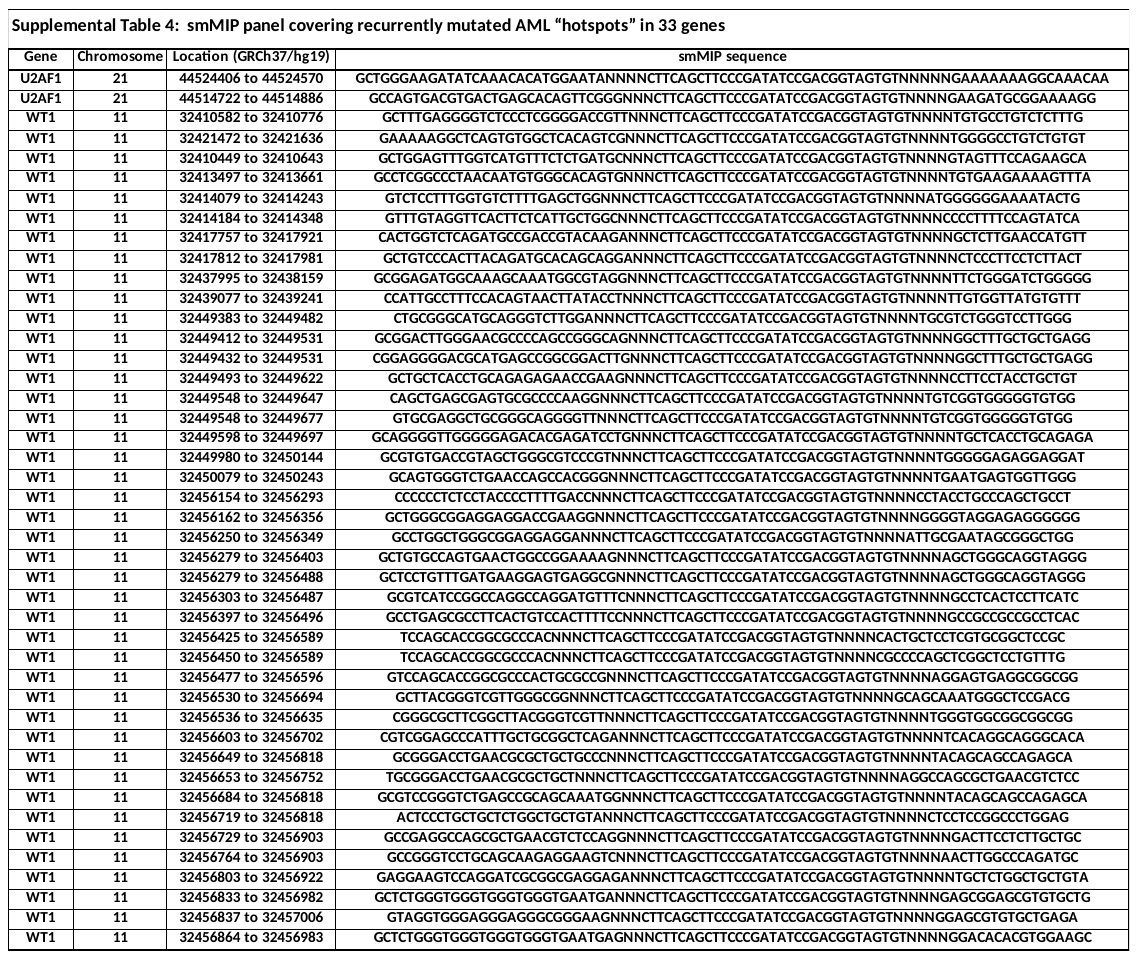

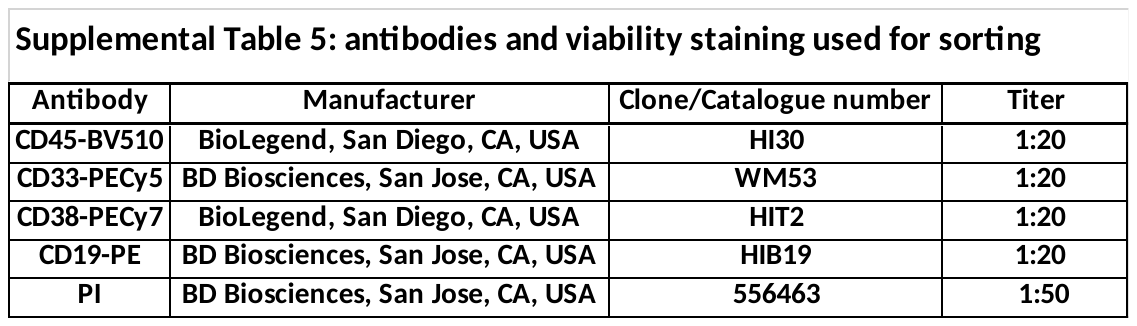

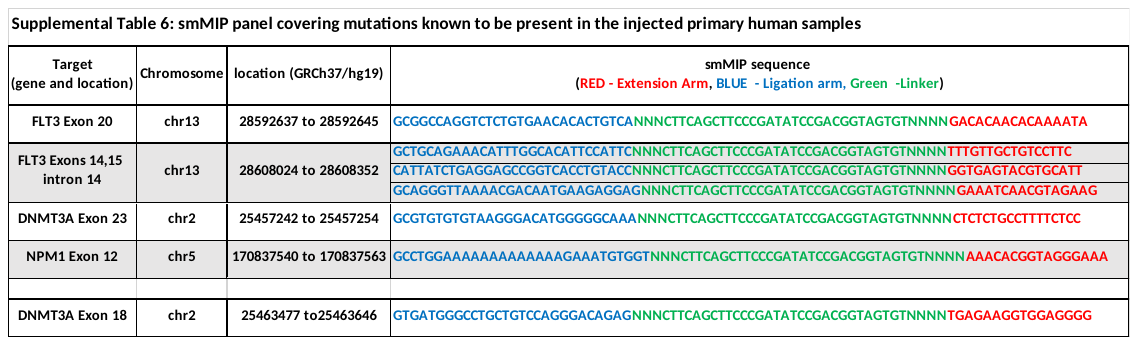

**References**

1. Shlush LI, Zandi S, Mitchell A, et al. Identification of pre-leukaemic haematopoietic stem cells in acute leukaemia.

Nature. 2014; 506(7488):328-333.

2. Kotecha N, Krutzik PO, Irish JM. Web-based Analysis and Publication of Flow Cytometry Experiments.

Current Protocols in Cytometry. 2010; Chapter 10, Unit 10.17: 1-40.

3. Amir ED, Davis KL, Tadmor MD, et al. viSNE enables visualization of high dimensional single-cell data and reveals phenotypic heterogeneity of leukemia.

Nat Biotechnol. 2013; 31(6):545–552.

4. Hainaut P, Milner J. A structural role for metal ions in the "wild-type" conformation of the tumor suppressor protein p53.

Cancer Res. 1993; 53(8):1739–1742.

5. Tal P, Eizenberger S, Cohen E, et al. Cancer therapeutic approach based on conformational stabilization of mutant P53 protein by small peptides.

Oncotarget. 2016; 7(11):11817-11837.

6. Hiatt JB, Pritchard CC, Salipante SJ, et al. Single molecule molecular inversion probes for targeted, high-accuracy detection of low-frequency variation.

Genome Res. 2013; 23(5):843-854.

7. Boyle EA, O’Roak BJ, Martin BK, et al. MIPgen: optimized modeling and design of molecular inversion probes for targeted resequencing.

Bioinformatics. 2014; 30(18):2670-2672.

8. Bushnell B, Rood J, Singer E. BBMerge - Accurate paired shotgun read merging via overlap.

PLoS One. 2017; 12, e0185056.

9. Martin M. Cutadapt removes adapter sequences from high-throughput sequencing reads.

EMBnet Journal. 2011; 17(1):10-12. doi: <https://doi.org/10.14806/ej.17.1.200>

10. Li H. Aligning sequence reads, clone sequences and assembly contigs with BWA-MEM.

arXiv. 2013; 1303.3997.

11. Li H, Handsaker B, Wysoker, A, et al. The Sequence Alignment/Map format and SAMtools.

Bioinformatics. 2009; 25: 2078-2079.

12. McKenna A, Hanna M, Banks E, et al. The Genome Analysis Toolkit: a MapReduce framework for analyzing next-generation DNA sequencing data.

Genome Res. 2010; 20:1297-303.

13. Koboldt DC, Zhang Q, Larson DE, et al. VarScan 2: somatic mutation and copy number alteration discovery in cancer by exome sequencing.

Genome Res. 2012; 22:568-576.

14. Rimmer A, Phan H, Mathieson I, et al. Integrating mapping-, assembly- and haplotype-based approaches for calling variants in clinical sequencing applications.

Nat Genet. 2014; 46:912-918.

15. Wang K, Li M, Hakonarson H. ANNOVAR: functional annotation of genetic variants from high-throughput sequencing data.
    Nucleic Acids Res. 2010; 38:e164.
16. Butler A, Hoffman P, Smibert P, et al. Integrating single-cell transcriptomic data across different conditions, technologies, and species.

Nat Biotechnol. 2018; 36(5):411–420.

17. Boyle EI, Weng S, Gollub J, et al. GO:TermFinder–open source software for accessing gene ontology information and finding significantly enriched gene ontology terms associated with a list of genes.

Bioinformatics. 2004; (18):3710–3715.

18. Franzén O, Gan LM, Björkegren JLM. PanglaoDB: a web server for exploration of mouse and human single-cell RNA sequencing data.

Database (Oxford). 2019; 2019: article ID baz046; doi:10.1093/database/baz046.

19. Kachitvichyanukul V, Schmeiser B. Computer generation of hypergeometric random variates.

J Stat Comput Simul. 1985; 22:127–145.

20. Benjamini Y, Hochberg Y. Controlling the false discovery rate: a practical and powerful approach to multiple testing.

J R Stat Soc Ser B (Methodological). 1995; 57(1):289–300.

21. Chen EY, Tan CM, Kou Y, et al. Enrichr: interactive and collaborative HTML5 gene list enrichment analysis tool.

BMC Bioinformatics. 2013; 14:128.

22. Kuleshov MV, Jones MR, Rouillard AD, et al. Enrichr: a comprehensive gene set enrichment analysis web server 2016 update.

Nucleic Acids Res. 2016; 44(W1):W90-W97. doi: 10.1093/nar/gkw377.
